# Impairment of the glial phagolysosomal system drives prion-like propagation in a *Drosophila* model of Huntington’s disease

**DOI:** 10.1101/2023.10.04.560952

**Authors:** Graham H. Davis, Aprem Zaya, Margaret M. Panning Pearce

## Abstract

Protein misfolding, aggregation, and spread through the brain are primary drivers of neurodegenerative diseases pathogenesis. Phagocytic glia are responsible for regulating the load of pathogenic protein aggregates in the brain, but emerging evidence suggests that glia may also act as vectors for aggregate spread. Accumulation of protein aggregates could compromise the ability of glia to eliminate toxic materials from the brain by disrupting efficient degradation in the phagolysosomal system. A better understanding of phagocytic glial cell deficiencies in the disease state could help to identify novel therapeutic targets for multiple neurological disorders. Here, we report that mutant huntingtin (mHTT) aggregates impair glial responsiveness to injury and capacity to degrade neuronal debris in male and female adult *Drosophila* expressing the gene that causes Huntington’s disease (HD). mHTT aggregate formation in neurons impairs engulfment and clearance of injured axons and causes accumulation of phagolysosomes in glia. Neuronal mHTT expression induces upregulation of key innate immunity and phagocytic genes, some of which were found to regulate mHTT aggregate burden in the brain. Finally, a forward genetic screen revealed Rab10 as a novel component of Draper-dependent phagocytosis that regulates mHTT aggregate transmission from neurons to glia. These data suggest that glial phagocytic defects enable engulfed mHTT aggregates to evade lysosomal degradation and acquire prion-like characteristics. Together, our findings reveal new mechanisms that enhance our understanding of the beneficial and potentially harmful effects of phagocytic glia in HD and potentially other neurodegenerative diseases.

**SIGNIFICANCE STATEMENT:** Deposition of amyloid aggregates is strongly associated with neurodegenerative disease progression and neuronal cell loss. Many studies point to glial cells as dynamic mediators of disease, capable of phagocytosing toxic materials, but also promoting chronic inflammation and proteopathic aggregate spread. Thus, glia have emerged as promising therapeutic targets for disease intervention. Here, we demonstrate in a *Drosophila* model of Huntington’s disease that neuronal mHTT aggregates interfere with glial phagocytic engulfment, phagolysosomal processing, and innate immunity transcriptional responses. We also identify Rab10 as a novel modifier of prion-like transmission of mHTT aggregates. Our findings add to a growing narrative of glia as double-edged players in neurodegenerative diseases.

## INTRODUCTION

Neuron-glia crosstalk is critical for maintaining homeostasis in the central nervous system (CNS), and disruption of these intercellular interactions is increasingly recognized as a central component of many neurological disorders, including neurodegenerative diseases. Glia perform immune surveillance functions in the CNS and respond to neuronal injury by altering gene expression (Magaki et al., 2018) and clearing damaged cells (Raiders et al., 2021; Zheng and Tuszynski, 2023). These glial responses may initially be neuroprotective, but prolonged glial reactivity propels the neurodegenerative disease state, for example, by driving premature loss of living neurons or functional synapses (Neniskyte et al., 2011; Hong et al., 2016; Dejanovic et al., 2022) and inducing neuroinflammation (Liddelow et al., 2017). Expanding our understanding of how glia influence neuron function and survival could reveal promising new therapeutic strategies for neurodegenerative diseases.

A pathological hallmark of most neurodegenerative diseases is the accumulation of misfolded proteins into intra- or extracellular amyloid aggregates in vulnerable regions of the CNS (Knowles et al., 2014). Protein aggregates form due to age-associated decline in cellular protein folding capacity (Santra et al., 2019; Stein et al., 2022) and overwhelming of degradative pathways, including the ubiquitin-proteasome system, autophagy, and phagocytosis (Aman et al., 2021; Duong et al., 2021; Wodrich et al., 2022). As professional phagocytes of the brain, microglia and astrocytes clear damaged and dysfunctional cells (Paolicelli et al., 2011; Wakida et al., 2018; Herzog et al., 2019; Lee et al., 2021) and other pathogenic material, such as protein aggregates (Liu et al., 2017; Choi et al., 2020).

Defective glial clearance of debris may be a driving force underlying neurodegenerative disease, highlighted by a growing list of genetic risk variants associated with phagocytic and endolysosomal pathways (Podleśny-Drabiniok et al., 2020). Endolysosomal impairment promotes protein aggregate accumulation; in turn, aggregates can drive further endolysosomal dysfunction, including deacidification and membrane permeabilization of intracellular vesicles (Krasemann et al., 2017; Heckmann et al., 2019; Burbidge et al., 2022; Lee et al., 2022). Aggregates that escape degradation may gain the ability to spread and seed soluble proteins in a prion-like manner (Jucker and Walker, 2018; Monaco and Fraldi, 2020).

Mechanisms by which pathogenic aggregates dysregulate endolysosomal processing remain largely unknown. Intracellular membrane fusion events that regulate endo/phagosome maturation are catalyzed by Rab GTPases (Ng and Tang, 2008; Chan et al., 2011; Langemeyer et al., 2018), enzymes that cycle between active (GTP-bound) and inactive (GDP-bound) states to organize endomembranes into distinct functional domains (Hall, 1990; Chan et al., 2011). Rab dysfunction is implicated in several neurodegenerative diseases—e.g., upregulation of Rab4, Rab5, Rab7, and Rab27 has been observed in Alzheimer’s disease (AD) (Ginsberg et al., 2011), and several Rabs are known substrates of leucine rich repeat kinase 2 (LRRK2), a genetic risk factor in familial Parkinson’s disease (PD) (Jeong et al., 2018). Intriguingly, spread of ɑ-synuclein between cultured neuronal and enteroendocrine cells is mediated by the LRRK2 substrate Rab35 (Bae et al., 2018; Rodrigues et al., 2022), suggesting that Rab-dependent processes may contribute to formation and propagation of pathogenic aggregate seeds.

Here, we investigated the impact of protein aggregates generated in neurons on phagocytic glial cell functions in a *Drosophila* model of Huntington’s disease (HD). We report that neuronal expression of aggregation-prone mutant huntingtin (mHTT) protein reduces the ability of glia to clear axonal debris and to mount phagocytic and innate immunity transcriptional responses following acute nerve injury.

We also observed mHTT-induced changes to numbers of glial lysosomes and Rab+ vesicles in uninjured brains, and identify Rab10 as a novel modifier of prion-like spreading of mHTT aggregates in adult fly brains. Together, these studies shed light on mechanisms by which phagocytic glia respond to and are impaired by accumulation of pathogenic aggregates in neurons.

## MATERIALS AND METHODS

### Fly husbandry

Fly stocks and crosses were raised on standard cornmeal/molasses media on a 12 hr light/12 hr dark cycle at 25°C, unless otherwise noted. No sex-specific differences were observed in any experiments, so both males and females were utilized in this study. Transgenic or mutant flies were either generated for this study (described below), purchased from Bloomington Drosophila Stock Center, or kindly provided by collaborators. Genotypes and sources of all fly stocks used in this study are listed in Table 1, and complete genotypes of flies used in each figure are listed in Table 2.

**Table 1.**
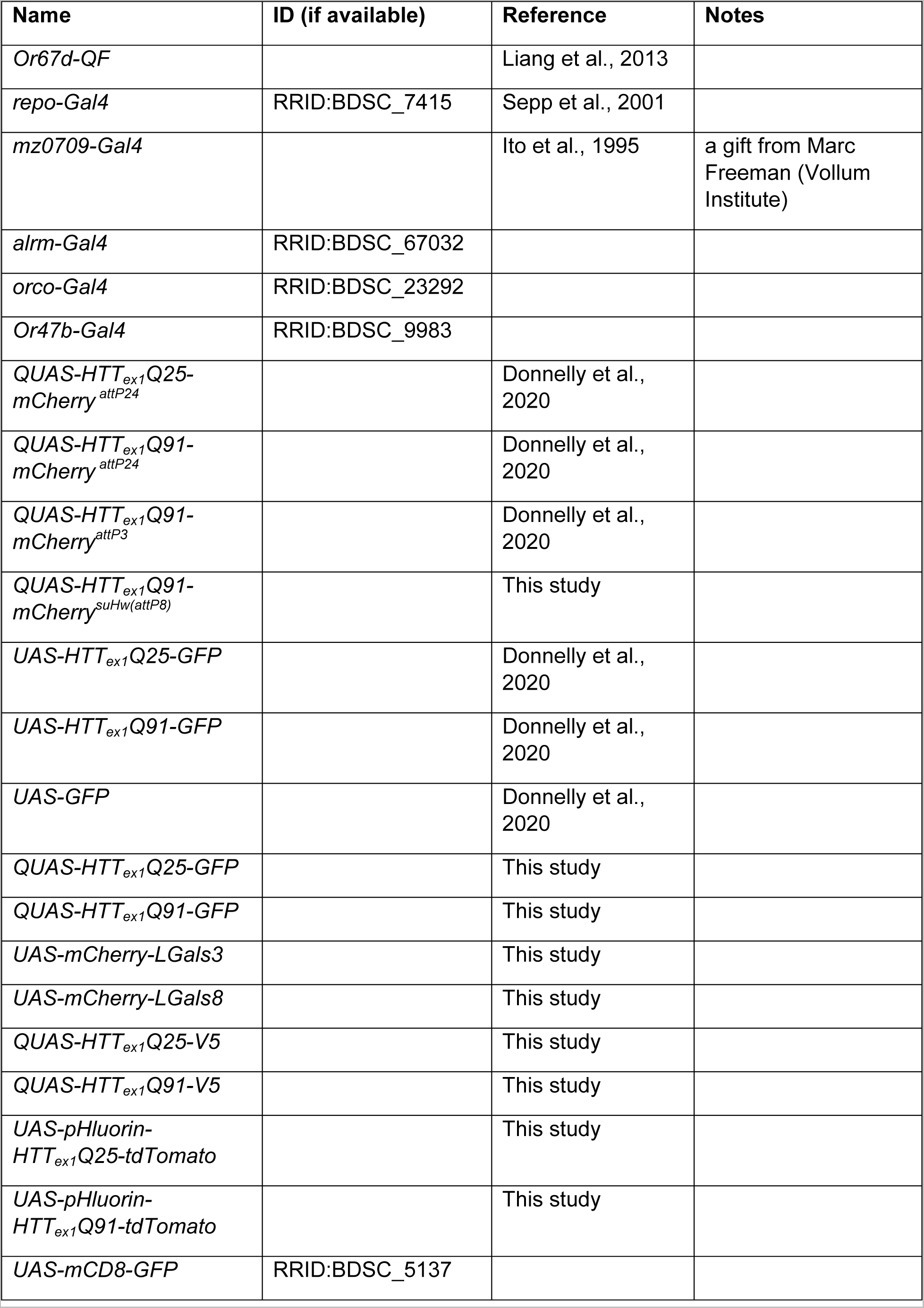
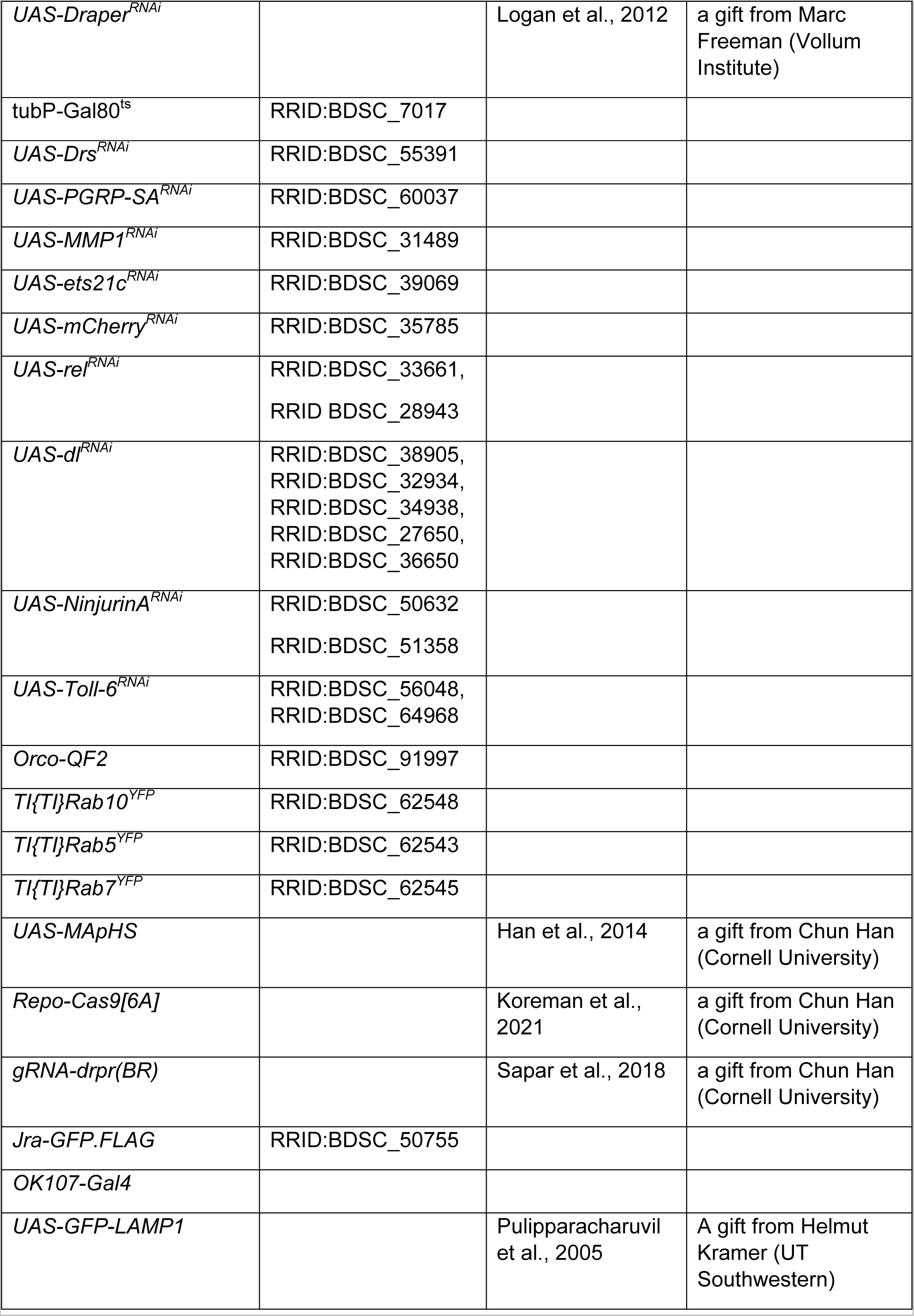
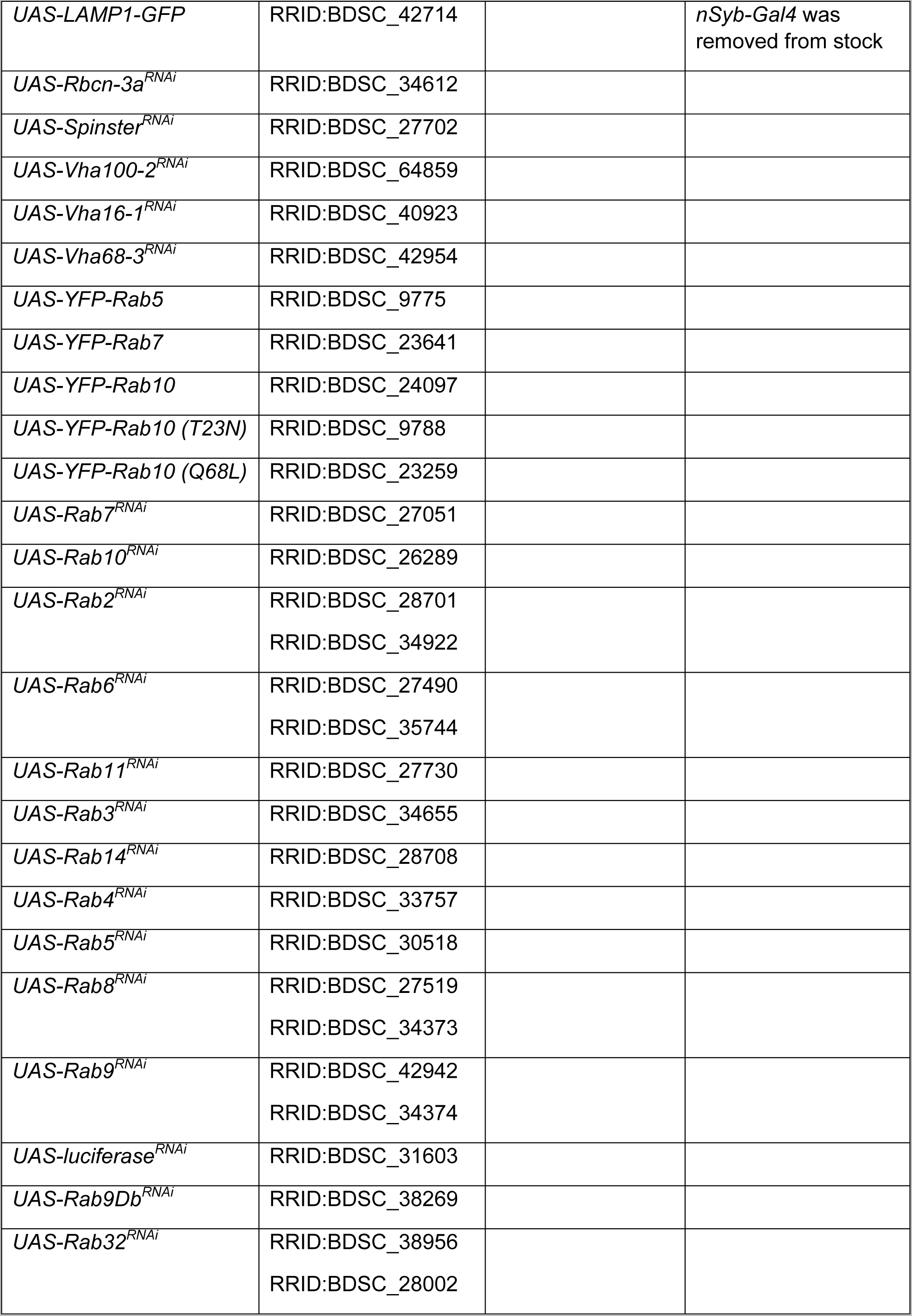
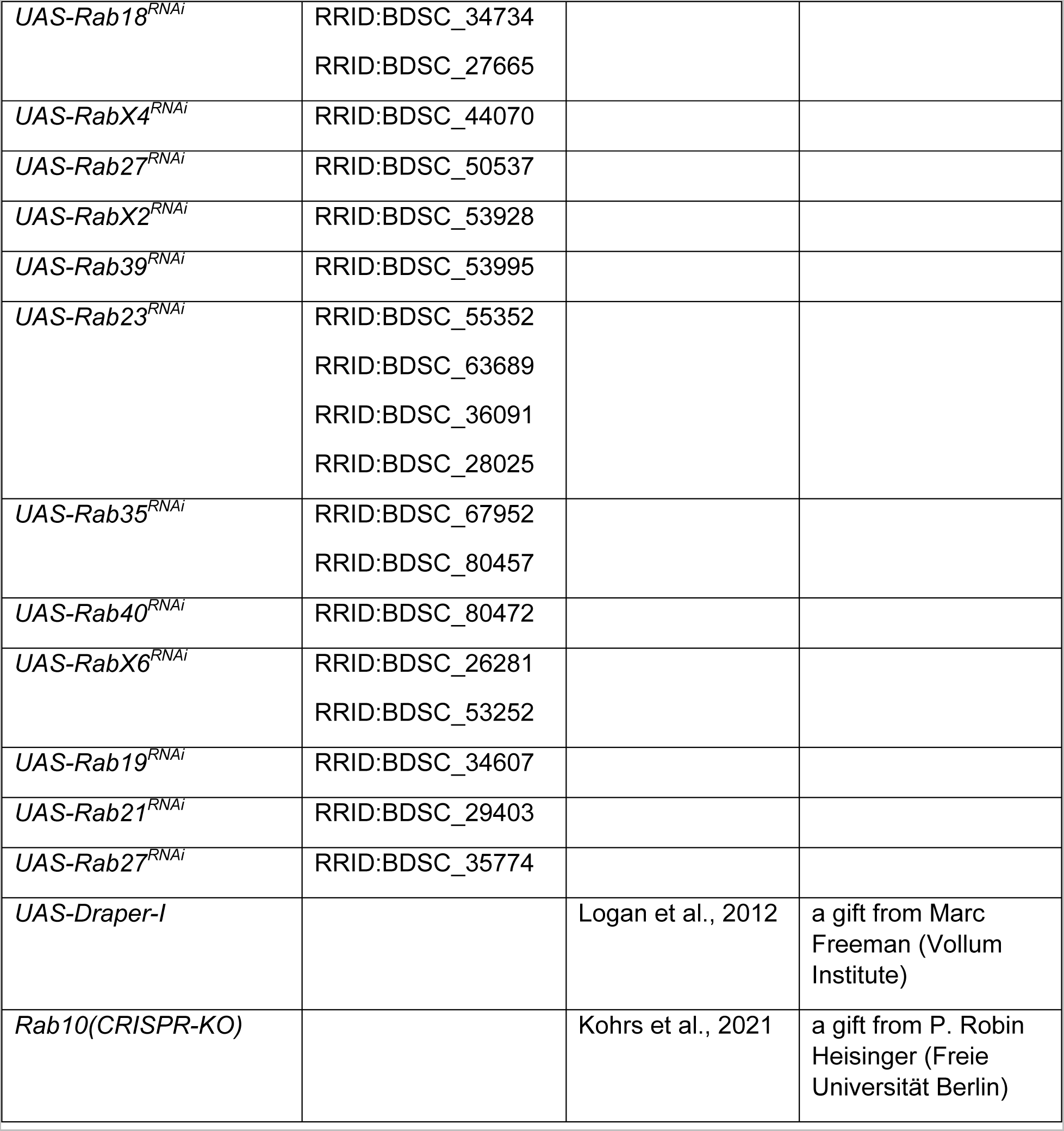
*Drosophila* genotype and source information.

**Table 2.**
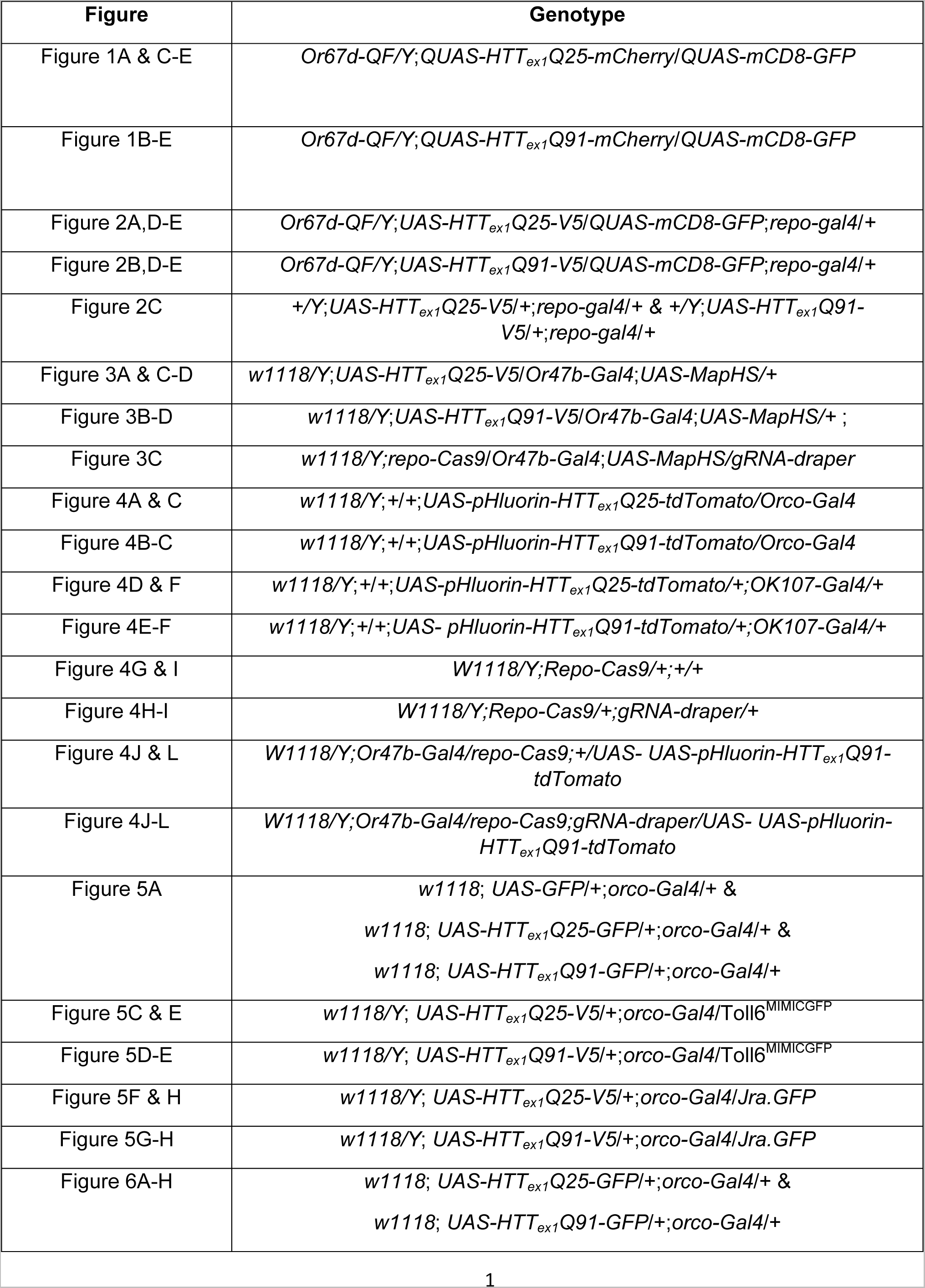
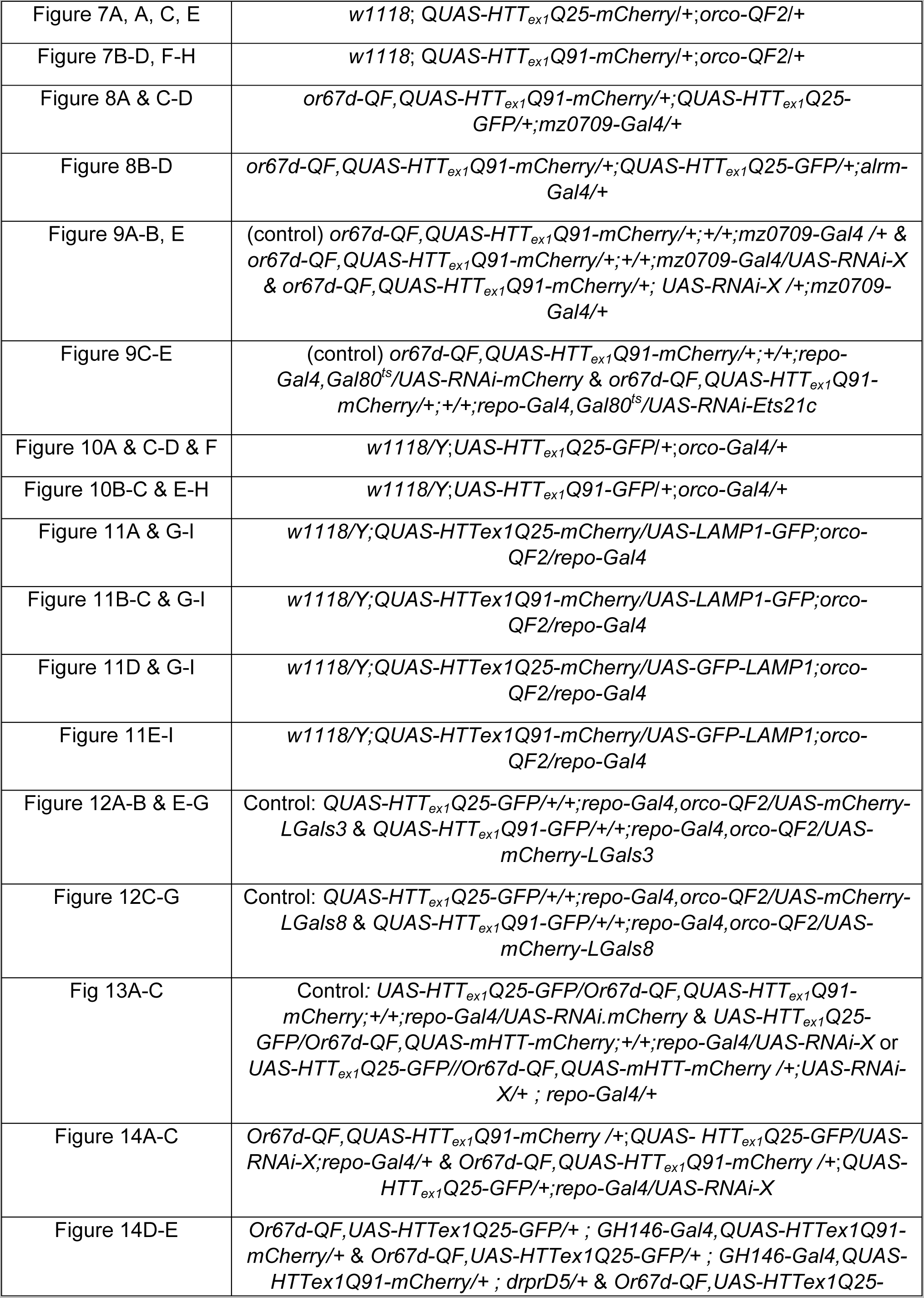
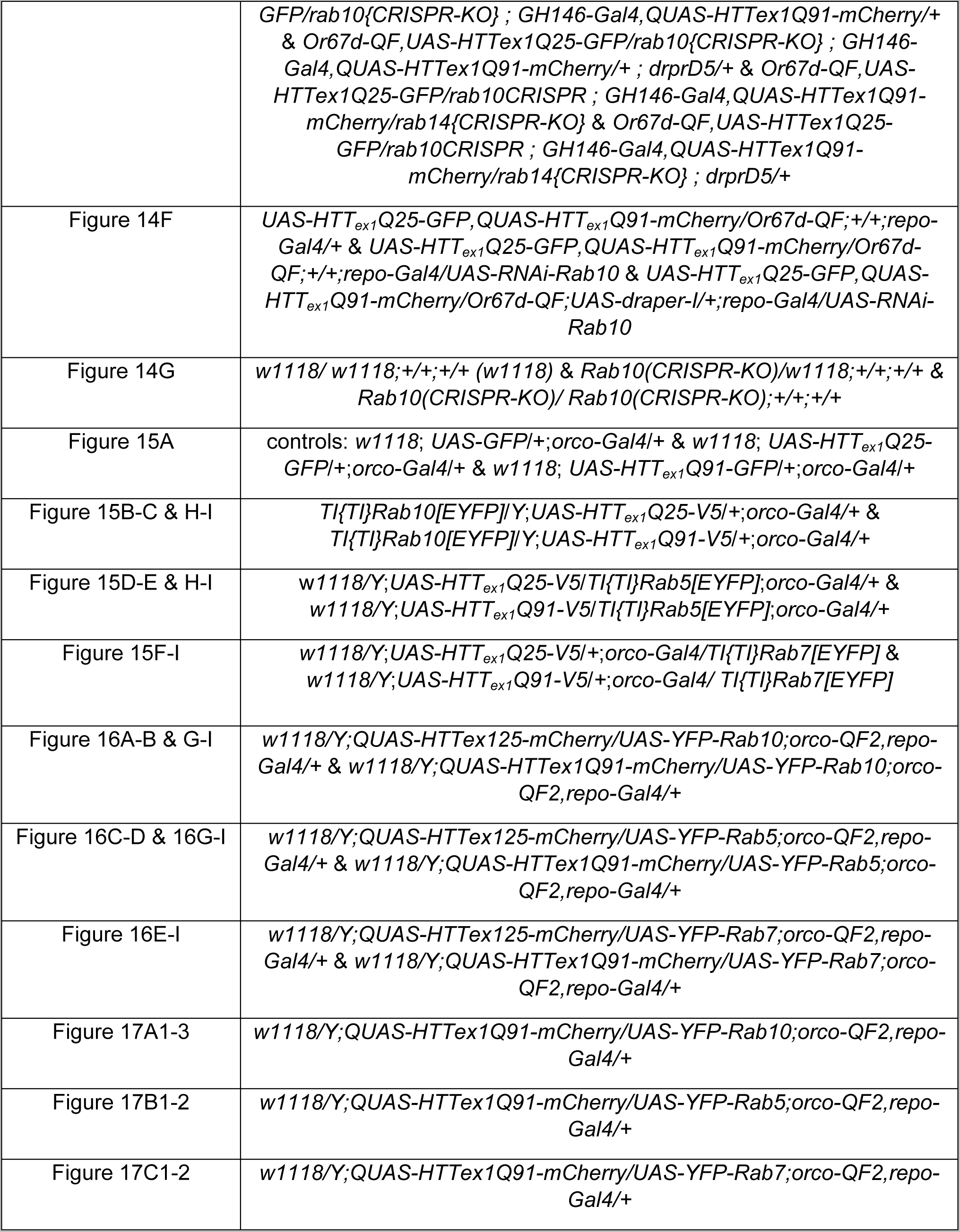
Genotypes of flies used in all figures.

Acute antennal nerve injury was performed by bilateral removal of the second and third antennal segments from anesthetized adult flies (MacDonald et al., 2006). For quantitative PCR analyses, maxillary palps were removed in addition to third antennal segments to sever all olfactory receptor neuron (ORN) axons. Axotomized flies were incubated for the indicated times on standard media prior to dissection and processing for imaging.

### Cloning and Drosophila transgenesis

pUASTattB(HTT_ex1_Q25-V5) and pUASTattB(HTT_ex1_Q91-V5) plasmids were cloned by PCR amplification of HTT exon 1 (HTT_ex1_) cDNA using a reverse primer that inserted an in-frame, C-terminal V5 epitope tag (GKPIPNPLLGLDST) and ligation into the pQUASTattB vector backbone via XhoI and XbaI restriction sites. pQUASTattB(HTT_ex1_Q25-GFP) and pQUASTattB(HTT_ex1_Q91-GFP) were generated by replacing the mCherry sequence in pQUASTattB(HTT_ex1_Q25-mCherry) and pQUASTattB(HTT_ex1_Q91-mCherry) plasmids previously generated by our lab (Pearce et al., 2015) with an in-frame, C-terminal GFP sequence via Gibson Assembly (New England Biolabs, Inc. Ipswich, MA). pUASTattB(pHluorin-HTT_ex1_Q25-tdTomato) and pUASTattB(pHluorin-HTT_ex1_Q91-tdTomato) plasmids were generated by PCR amplification of HTT_ex1_ cDNAs and subcloning to replace the CD4 sequence in the pUASTattB(MApHS) plasmid (Han et al., 2014) via In-Fusion cloning (Takara Bio USA, Inc., Mountain View, CA). pUASTattB(mCherry-Galectin-3) and pUASTattB(mCherry-Galectin-8) were cloned by PCR amplification of human LGals3 and LGals8 cDNAs from the pHAGE-mKeima-LGALS3 and pHAGE-FLAG-APEX2-LGALS8 plasmids (Addgene plasmids #175780 and #175758, Watertown, MA) (Eapen et al., 2021) and insertion downstream of an in-frame mCherry sequence in the pUASTattB vector backbone via Gibson Assembly.

Plasmids were microinjected into *w-* embryos with the su(Hw)attP8, attP24, VK19, or VK27 φC31 attP integration sites at BestGene, Inc (Chino Hills, CA). Table 1 lists detailed genotype information for all transgenic flies generated in this study.

### Drosophila brain dissection and sample preparation

Adult fly brains were dissected in ice-cold phosphate-buffered saline (PBS) containing either no detergent or 0.03% (PBS/0.03T), 0.1% (PBS/0.1T), or 0.3% (PBS/0.3T) Triton X-100. Dissected brains were fixed in 4% paraformaldehyde (PFA) in the dark at room temperature (RT) for 20 minutes. For imaging of GFP or mCherry fluorescence signals, brains were washed 7X in PBS/0.03T buffer before incubation in Slowfade Gold Antifade Mountant (Invitrogen, Carlsbad, CA). When imaging pHluorin-tagged and other pH-sensitive constructs, such as GFP-LAMP1, brains were washed 7X in PBS for at least 50 minutes at RT before incubation in Slowfade. For immunostaining, brains were washed 7X in PBS/0.3T after fixation, blocked in PBS/0.3T containing 5% normal goat serum (Lampire Biological Laboratories, Pipersville, PA) for 30 min at RT, incubated in primary antibodies diluted in blocking solution for 24-48 hours at 4°C, washed 7X in PBS/0.3T, and then incubated in secondary antibodies diluted in blocking solution for 16-20 hours at 4°C. After a final set of 7X PBS/0.3T washes, dissected brains were incubated in Slowfade. For Magic Red and LysoTracker staining, brains were dissected in PBS and incubated in 1:1,000 LysoTracker Red DND-99 (Invitrogen, Carlsbad, CA) or 1:1,250 Magic Red (ImmunoChemistry, Davis, CA) diluted in PBS for 20 minutes at RT. Brains were washed 5X in PBS for 15 min total, fixed in 4% PFA in PBS/0.1T for 20 minutes at RT, washed 6X in PBS/0.1T for 30 min total, and incubated in Slowfade. Following incubation in Slowfade for 1-24 hours at 4°C in the dark, all brains were bridge-mounted in Slowfade on glass microscopy slides under #1.5 cover glass, and edges were sealed using clear nail polish.

Primary antibodies used in this study include: chicken anti-GFP (RRID: AB_300798; 1:1,000; Abcam, Cambridge, UK), chicken anti-GFP (RRID: AB_2534023; 1:500; Thermo Fisher Scientific Inc., Waltham, MA), rat anti-N-cadherin (clone DN-Ex #8; RRID: AB_528121; 1:75; Developmental Studies Hybridoma Bank, Iowa City, IA), mouse anti-Repo (clone 8D12; RRID: AB_528448; 1:25; Developmental Studies Hybridoma Bank, Iowa City, IA), mouse anti-V5 (RRID: AB_2556564; 1:125; Thermo Fisher Scientific Inc., Waltham, MA), and rabbit anti-Draper (1:500; a kind gift from Marc Freeman, Vollum Institute, Portland, OR). Secondary antibodies include: AlexaFluor 405 goat anti-rabbit (RRID: AB_221605; 1:250), FITC-conjugated donkey anti-chicken (RRID: AB_2340356; 1:250; Jackson Immuno Research Labs, West Grove, PA), AlexaFluor 568 goat anti-mouse (RRID: AB_2534072; 1:250), AlexaFluor 647 goat anti-rabbit (RRID: AB_2535812; 1:250), and AlexaFluor 647 goat anti-rat IgGs (RRID: AB_141778; 1:250) (Invitrogen, Carlsbad, CA).

### Image acquisition

All microscopy data were collected on a Leica SP8 laser-scanning confocal system equipped with 405 nm, 488 nm, 561 nm, and 633 nm lasers and 40X 1.3 NA or 63X 1.4 NA oil objective lenses. Leica LAS-X software was used to establish optimal settings during each microscopy session and to collect optical z-slices of whole-mounted brain samples with Nyquist-optimized sampling criteria.

Optical zoom was used to magnify and establish a region of interest (ROI) in each sample. For images showing a single glomerulus, confocal slices were collected to generate ∼73 x 73 x 20 μm (*xyz*) stacks, with z-axis boundaries established using fluorescent signal in DA1 or VA1lm ORN axons. For images of a single antennal lobe, confocal slices were collected to generate ∼117 x 117 x 26 μm (*xyz*) stacks, which were located using HTT_ex1_ fluorescence in ORN axons.

### Post-imaging analysis

Raw confocal data were analyzed in 2D using ImageJ/FIJI (RRID:SCR_002285; NIH, Bethesda, MD) or in 3D using Imaris image analysis software (RRID:SCR_007370; Bitplane, Zürich, Switzerland). Methods used for image segmentation and semi-automated quantification of fluorescent signals were previously described (Donnelly et al., 2020). Briefly, raw confocal data were cropped to establish an ROI for further analysis and displayed as a 2D sum intensity projection (ImageJ) or a 3D volume (Imaris). Background fluorescence was subtracted from raw confocal images. mCD8-GFP and HTT_ex1_-mCherry fluorescent signals was segmented using the ‘Surfaces’ function in Imaris (surface detail = 0.25 μm, background subtraction = 0.75 μm). Using the ‘split touching objects’ option, seed point diameter was set to 0.85 μm. pHluorin- and tdTomato-labeled VA1lm ORN axons were quantified in central 30 x 30 pixels, 50-slice ROIs from sum intensity projections of each VA1lm glomerulus. pHluorin- and tdTomato-labeled Or83b+ ORN axons or MBN soma were quantified from central 100 x 100 pixels, 75-slice ROIs of each antennal lobe or mushroom body calyx, and background intensity was subtracted from sum intensity projections. Quantification of Toll-6^MIMICGFP^ fluorescence was performed using the ‘Spots’ function in Imaris (*xy* diameter = 0.5 μm, *z*-diameter = 1.0 μm), and glial nuclei-associated GFP-Jra signal was quantified using the “Surfaces” function to identify Repo+/GFP+ nuclei (surface detail = 0.2 μm, background subtraction = 1.6 μm, number of voxels >10).

mCherry-tagged mHTT_ex1_ and GFP-tagged wtHTT_ex1_ aggregates were identified and quantified as previously reported (Donnelly et al., 2020). Briefly, mCherry+ surfaces were segmented and measured in Imaris (surface detail = 0.2 μm, background subtraction = 0.45 μm, seed point diameter = 0.85 μm). Seeded wtHTT_ex1_ aggregates were identified as mHTT_ex1_-mCherry+ objects that overlapped with wtHTT_ex1_-GFP signal. Intracellular vesicles were identified using the ‘Surfaces’ algorithm in Imaris (surface detail = 0.2 μm, background subtraction = 0.4 μm, volume >0.001 μm^3^) to segment fluorescent signals associated with lysosomes (MR+, LTR+, LAMP1-GFP+, GFP-LAMP1+, mCherry-Galectin-3+, or mCherry-Galectin-8+) or phagosomes (YRab+ or YFP-Rab+). Intracellular vesicles associated with mHTT_ex1_ aggregates were identified by filtering for lysosomal or phagosomal surfaces within 0.2 μm of mHTT_ex1_ objects using the “Shortest Distance” calculation in Imaris.

### qPCR

Transgenic *Drosophila* were flash frozen in liquid nitrogen, and heads were collected using a microsieve with a 230 nm filter to separate bodies from heads and 170 nm filter to separate heads from appendages. Total RNA was extracted from isolated fly heads using the Zymo Direct-Zol RNA miniprep kit (Zymo Research, Irvine, CA). Extracted RNA was quantified on a Nanodrop 2000 (Thermo Fisher Scientific), and equal quantities of each sample were subjected to cDNA synthesis using the High-Capacity cDNA Reverse Transcription Kit (Applied Biosystems, Waltham, MA). qPCR was performed on a T100 Thermal Cycler with a CFX384 Real-Time System (Bio-Rad Laboratories, Hercules, CA) using 10 ng of input RNA per replicate and TB Green Premix Ex Taq II (Takara Bio, Kusatsu, Japan). Sequences of all qPCR primers used in this study are listed in Table 3.

**Table 3.**
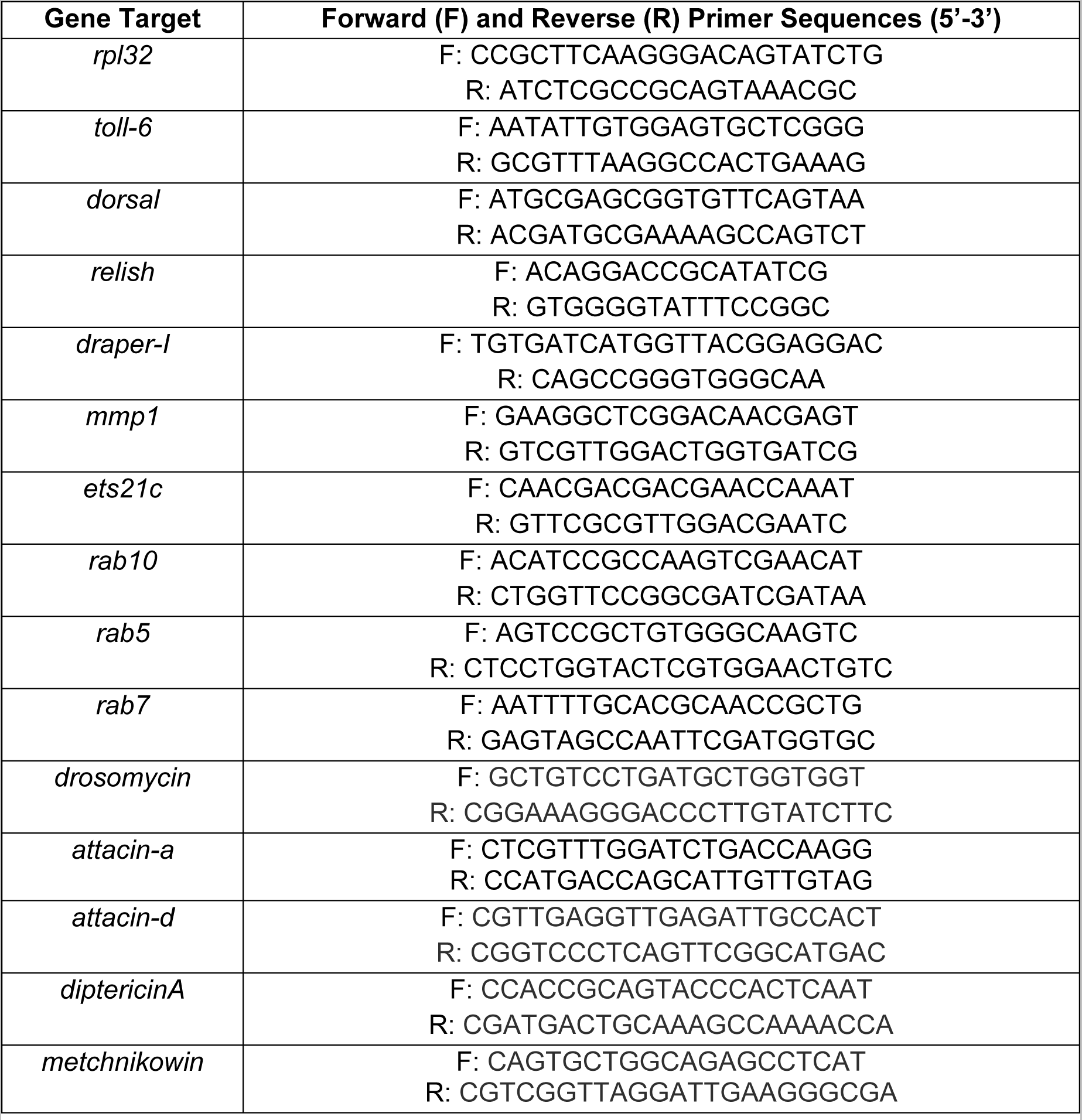
Primer sequences used for qPCR analyses.

### Experimental design and statistical analyses

All quantified data were organized and analyzed in Prism 9 software (Graphpad, San Diego, CA). Power analyses to determine appropriate sample sizes for each experiment were calculated using a Sample Size calculator available at ClinCalc.com (ɑ = 0.05, β = 0.2). All quantifications are graphed as mean ± standard error of the mean (s.e.m). A single glomerulus or antennal lobe represented one biological replicate. Statistical comparisons were performed using the following tests where appropriate: unpaired, two-tailed *t*-test when comparing two samples and ANOVA followed by *post hoc* multiple comparison tests when comparing ≥3 samples. Detailed statistical information for each experiment, including sample sizes (*n*), means, and test results are summarized in Table 4.

## RESULTS

### Expression of mHTT exon 1 fragments in neurons inhibits phagocytic clearance of axonal debris

HD is a monogenic neurodegenerative disease caused by expansion of a CAG repeat region in exon 1 of the *huntingtin* (*HTT/IT15)* gene beyond a pathogenic threshold of 37 repeats, leading to production of mHTT proteins containing expanded N-terminal polyglutamine (polyQ≥37) tracts (Scherzinger et al., 1999). mHTT proteins are prone to misfolding and accumulate into insoluble aggregates (Wanker, 2000), whereas wild-type HTT (wtHTT) proteins containing polyQ≤36 tracts remain soluble unless seeded by a pre-formed HTT aggregate (Preisinger et al., 1999; Chen et al., 2001). To recapitulate these molecular features of HD, we generated transgenic flies that express N-terminal exon 1 fragments of human mHTT (mHTT_ex1_) containing a polyQ91 tract or wtHTT (wtHTT_ex1_) with a polyQ25 tract via the GAL4-UAS (Brand and Perrimon, N., 1993) or QF-QUAS binary expression systems (Potter and Luo, 2011). As we have previously reported (Pearce et al., 2015; Donnelly et al., 2020), fluorescent protein fusions of mHTT_ex1_ and wtHTT_ex1_ appeared punctate (i.e., aggregated or insoluble) or diffuse (i.e., soluble), respectively, in axon termini of the DA1 type of olfactory receptor neurons (ORNs), which synapse in the DA1 glomerulus of the antennal lobe in the central brain of adult flies (Couto et al., 2005) (Fig. 1A and B).

**Figure 1.**
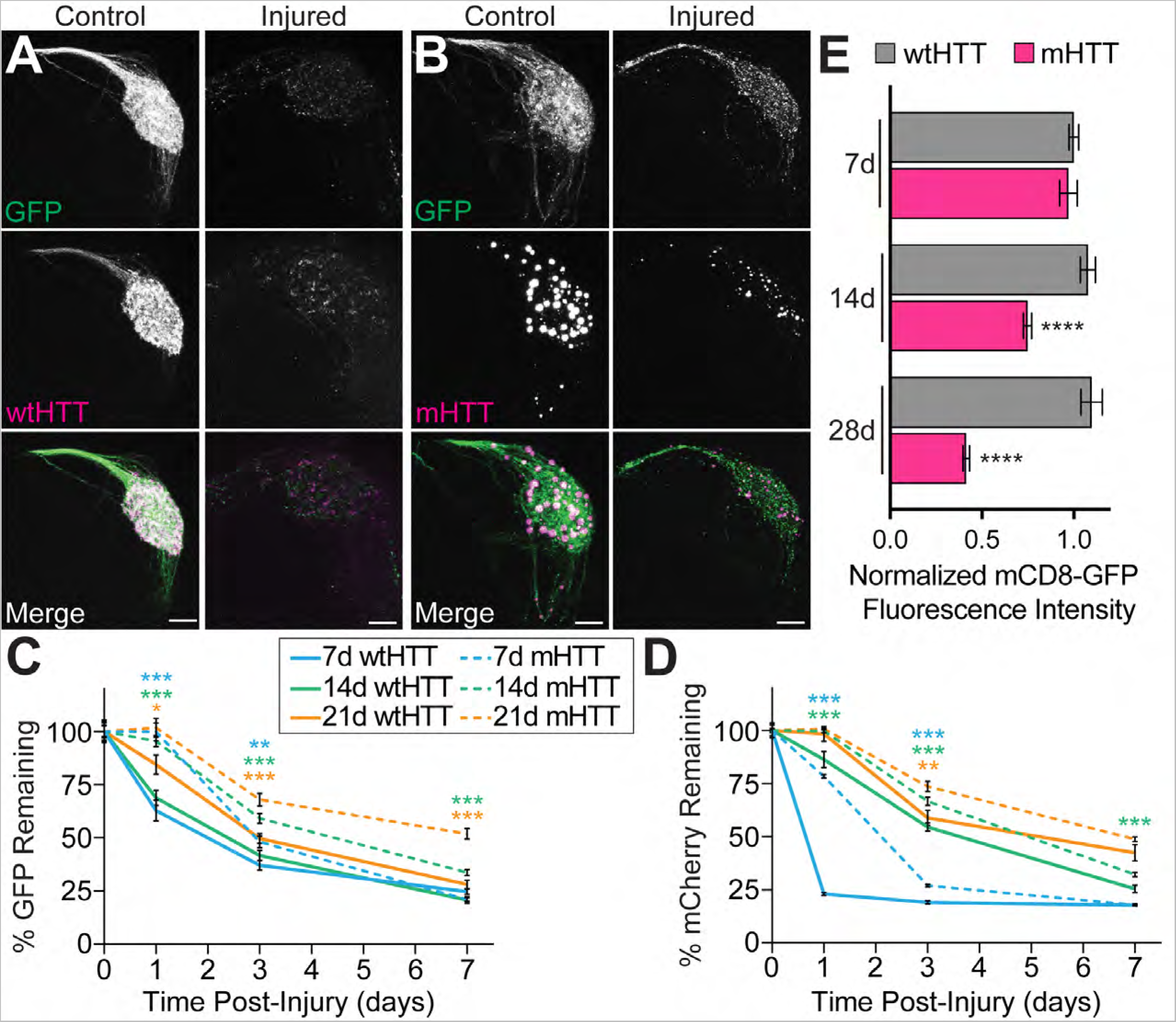
mHTT_ex1_ expression in ORNs impairs clearance of injured axons. (**A-B**) Maximum intensity projections of mCD8-GFP-labeled DA1 ORN axons expressing (**A**) HTT_ex1_Q25-(wtHTT_ex1_) or (**B**) HTT_ex1_Q91-mCherry (mHTT_ex1_) in 7 day-old uninjured flies (*left*) or 14 day-old flies subjected to bilateral antennal nerve axotomy 7 days earlier (*right*). Scale bars = 5 μm. (**C-D**) Quantification of (**C**) mCD8-GFP+ and (**D**) HTT_ex1_-mCherry+ DA1 ORN axons remaining in 7, 14, and 28 day-old uninjured flies or flies at 1, 3, and 5 days post-injury. (**E**) Quantification of mCD8-GFP+ DA1 ORN axons in 7, 14, and 28 day-old flies expressing HTT_ex1_Q25- or HTT_ex1_Q91-mCherry in DA1 ORNs. All quantified data were normalized to uninjured 1 day-old adults and graphed as mean ± s.e.m.; \**p*<0.05, \*\**p*<0.01, \*\*\**p*<0.001, \*\*\*\**p*<0.0001 by unpaired two-tailed *t*-test.

To monitor the capacity of glia to maintain CNS homeostasis in the presence of mHTT_ex1_ aggregates, we performed a series of experiments that quantified glial responses to acute injury in the adult fly CNS. Surgical removal of the second and third antennal segments initiates Wallerian degeneration of ORN axons, inducing a robust phagocytic glial response that involves upregulation of the scavenger receptor, Draper, and clearance of axonal debris within 7 days (MacDonald et al., 2006). To determine if neuronal mHTT_ex1_ expression affects the ability of glial cells to efficiently clear axonal debris, we co-expressed mCherry-tagged HTT_ex1_ transgenes with membrane-targeted GFP (mCD8-GFP) in DA1 ORNs and measured GFP and mCherry fluorescence intensities following antennal nerve axotomy (Fig. 1A-D). Interestingly, mHTT_ex1_ expression was associated with reduced steady-state mCD8-GFP levels in DA1 ORN axons in 2 and 4 week-old flies (Fig. 1E), likely due to neurotoxicity caused by accumulation of mHTT_ex1_ aggregates over time. Quantification of DA1 ORN axons remaining after antennal nerve injury revealed that clearance of axonal debris was reduced in flies co-expressing neuronal mHTT_ex1_ compared with controls expressing wtHTT_ex1_ (Fig. 1C). This effect could be observed as early as 1 day post-injury, suggesting that mHTT_ex1_ causes defects in both clearance and engulfment of axonal debris. Delayed clearance of axonal debris was exacerbated in older (14 and 28 day-old) mHTT_ex1_-expressing flies (Fig. 1C), possibly related to a decline in Draper activity during normal aging (Purice et al., 2016) and/or enhanced glial dysfunction compounded by mHTT_ex1_ aggregate accumulation. Quantification of mCherry fluorescence indicated that clearance of axonal mHTT_ex1_ was also slowed compared to wtHTT_ex1_ (Fig. 1D), suggesting that glia are deficient in degrading both axonal debris and neuronal mHTT_ex1_ aggregates.

We also tested whether formation of mHTT_ex1_ aggregates in glial cells impacts ORN axonal debris clearance following acute injury. Restricting expression of mHTT_ex1_ to glia resulted in appearance of heterogeneously-sized mHTT_ex1_ aggregates throughout the brain (Fig. 2A). Glial mHTT_ex1_ aggregates also slowed injury-induced clearance of mCD8-GFP-labeled axons compared with wtHTT_ex1_-expressing controls (Fig. 2B-D), though glial mHTT_ex1_ expression did not affect DA1 ORN axon abundance in 1 day-old flies (Fig. 2E). Together, these findings indicate that mHTT_ex1_ aggregates originating in either neurons or glia slow efficient clearance of injured ORN axons by phagocytic glia.

**Figure 2.**
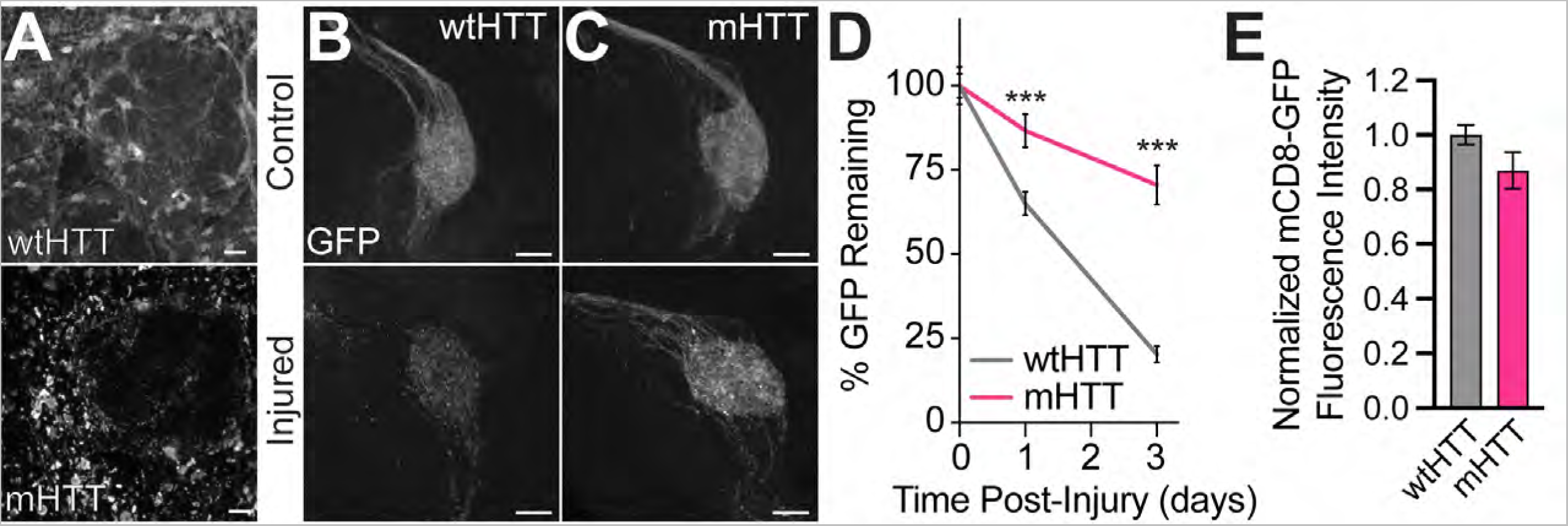
mHTT_ex1_ expression in glia is associated with reduced ORN axon clearance post-injury. (**A-B**) Maximum intensity projections of antennal lobes from 5-6 day-old flies expressing HTT_ex1_Q25-(*top*) or HTT_ex1_Q91-V5 (*bottom*) in glia and immunostained with anti-V5. Scale bars = 10 μm. (**B-C**) Maximum intensity projections of mCD8-GFP-labeled DA1 ORN axons in 1 day-old flies expressing (**B**) HTT_ex1_Q25- or (**C**) HTT_ex1_Q91-V5 in repo+ glia. Scale bars = 5 μm. (**D**) Quantification of mCD8-GFP+ DA1 ORN axons in flies expressing HTT_ex1_Q25- or HTT_ex1_Q91-V5 in repo+ glia, either uninjured or at 1 and 3 days post-injury. (**E**) Quantification of mCD8-GFP+ DA1 ORN axons in 1 day-old flies expressing HTT_ex1_Q25- or HTT_ex1_Q91-mCherry in repo+ glia. All data were normalized to uninjured 1 day-old adult flies and graphed as mean ± s.e.m.; \*\*\**p*<0.001 by unpaired two-tailed *t*-test.

### Neuronal mHTT aggregates impair nascent phagosome formation

Phagocytosis occurs via 4 major steps: (1) extension of phagocyte membranes toward extracellular “find me” cues, (2) recognition of “eat me” signals by scavenger receptors, (3) cytoskeletal and plasma membrane reorganization to surround and internalize extracellular material, and (4) maturation of nascent phagosomes through sequential endomembrane fusions that culminate at the lysosome (Vieira et al., 2002). We have previously reported that the conserved scavenger receptor, Draper/Ced-1/MEGF10 (MacDonald et al., 2006; Evans et al., 2015; Iram et al., 2016), regulates engulfment, clearance, and intercellular spreading of mHTT_ex1_ aggregates originating in ORN axons (Pearce et al., 2015; Donnelly et al., 2020). To determine whether delayed clearance of injured axons containing mHTT_ex1_ aggregates (Fig. 1C and 2D) could result from defective Draper-dependent engulfment, we co-expressed V5-tagged mHTT_ex1_ or wtHTT_ex1_ with the ratiometric membrane-associated pH sensor (MApHS), consisting of the transmembrane domain of CD4 flanked by N-terminal ecliptic pHluorin and C-terminal tdTomato (Han et al., 2014), in the VA1lm type of ORNs. Ecliptic pHluorin is brightest at pH 7.5 and dims as pH drops, quenching at pH <6.0 (Miesenböck et al., 1998), whereas tdTomato fluorescence is pH resistant. Thus, internalization of MApHS-labeled ORN axonal debris into a rapidly-acidified nascent phagosome can be monitored by calculating pHluorin:tdTomato fluorescence intensity ratios (Han et al., 2014). Indeed, pHluorin:tdTomato ratios in VA1lm axons were decreased at 14 and 25 hours post-axotomy (Fig. 3A-C), and this effect was lost when *draper* was deleted from glia using the tissue-specific CRISPR/Cas9-TriM method (Fig. 3C and E-G) (Poe et al., 2019), demonstrating that this construct accurately reports nascent phagosome acidification following engulfment. Notably, the decrease in pHluorin:tdTomato ratio following injury was less pronounced in VA1lm ORN axons co-expressing mHTT_ex1_ compared with wtHTT_ex1_-expressing controls (Fig. 3A-C), suggesting that mHTT_ex1_ aggregates impair nascent phagosome formation following engulfment.

**Figure 3.**
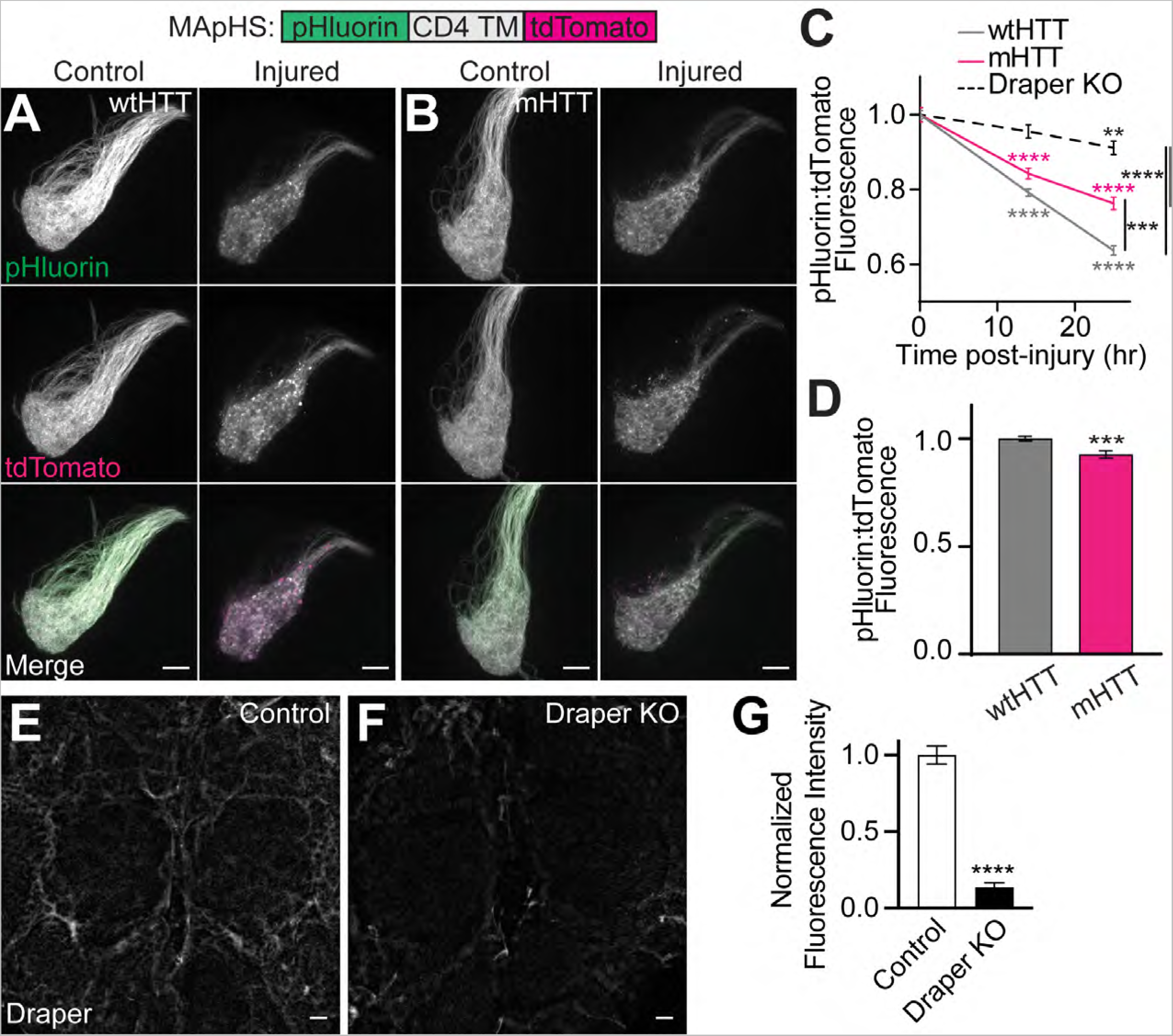
mHTT_ex1_ expression inhibits engulfment of injured ORN axons. (**A-B**) Maximum intensity projections of VA1lm ORN axons co-expressing the ratiometric phagocytic indicator, MApHS, and (**A**) HTT_ex1_Q25- or (**B**) HTT_ex1_Q91-V5 from 7 day-old uninjured flies (*left*) and flies 25 hours post-injury (*right*). Scale bars = 10μm. (**C**) pHluorin:tdTomato fluorescence intensity ratios calculated in VA1lm glomeruli from 7-day-old uninjured flies and flies at 14 or 25 hours post-injury. Data were normalized to the uninjured condition for each genotype. (**D**) pHluorin:tdTomato fluorescence intensity ratios in 7 day-old flies co-expressing MApHS with HTT_ex1_Q25- or HTT_ex1_Q91-V5 in VA1lm ORNs, normalized to wtHTT_ex1_ controls. Data are shown as mean ± s.e.m.; \*\**p*<0.01, \*\*\**p*<0.001, \*\*\*\**p*<0.0001 by unpaired two-tailed *t*-test. (**E-F**) Maximum intensity projections of the central brain from 6-7 day-old flies expressing (**E**) repo-Cas9 and (**F**) repo-Cas9 plus gRNAs targeting *draper* (“Draper KO”). Brains were immunostained with anti-Draper. Scale bars = 10 μm. (**G**) Quantification of Draper immunofluorescence in brains from flies shown in (**E-F**), normalized to control.

### Neuronal mHTT accumulates in low pH intracellular compartments

mHTT_ex1_ expression was associated with a slight but significant decrease in steady-state pHluorin:tdTomato ratios in MApHS-labeled axons compared with wtHTT_ex1_-expressing controls (Fig. 3D), suggesting that even in the absence of acute injury, mHTT_ex1_ signals for ORN axon engulfment. To test this, we generated transgenic flies that express mHTT_ex1_ or wtHTT_ex1_ fused to N-terminal pHluorin and C-terminal tdTomato fluorescent proteins, herein referred to as mHTT-associated pH sensor (mHApHS) or wtHTT-associated pH sensor (wtHApHS). Accumulation of HTT_ex1_ protein in low pH cellular compartments would cause a decrease in the ratio of pHluorin:tdTomato fluorescence for these constructs. wtHApHS and mHApHS transgenes were expressed in either Or83b+ ORNs, which encompass 70-80% of all adult ORNs (Fig. 4A-B) (Larsson et al., 2004), or mushroom body neurons (MBNs; Fig. 4D-E), which are downstream in the fly olfactory circuit and innervate the learning and memory center of the fly CNS (McGuire et al., 2001). In both ORN axons and MBN soma, pHluorin:tdTomato ratios associated with mHApHS were significantly decreased compared with wtHApHS controls (Fig. 4C and F), suggesting that mHTT_ex1_ proteins accumulate in acidified cellular compartments. To discriminate mHTT_ex1_ proteins engulfed by glia from mHTT_ex1_ internalized into neuronal autophagolysosomes, we measured VA1lm ORN axonal mHApHS-associated fluorescence in animals with glial *draper* loss-of-function (Fig. 4G-H). Glial *draper* knockout increased pHluorin:tdTomato ratios for axonal mHApHS (Fig. 4I), suggesting that at least some portion of mHTT_ex1_ aggregates accumulate in acidic portions of the glial phagolysosomal system.

**Figure 4.**
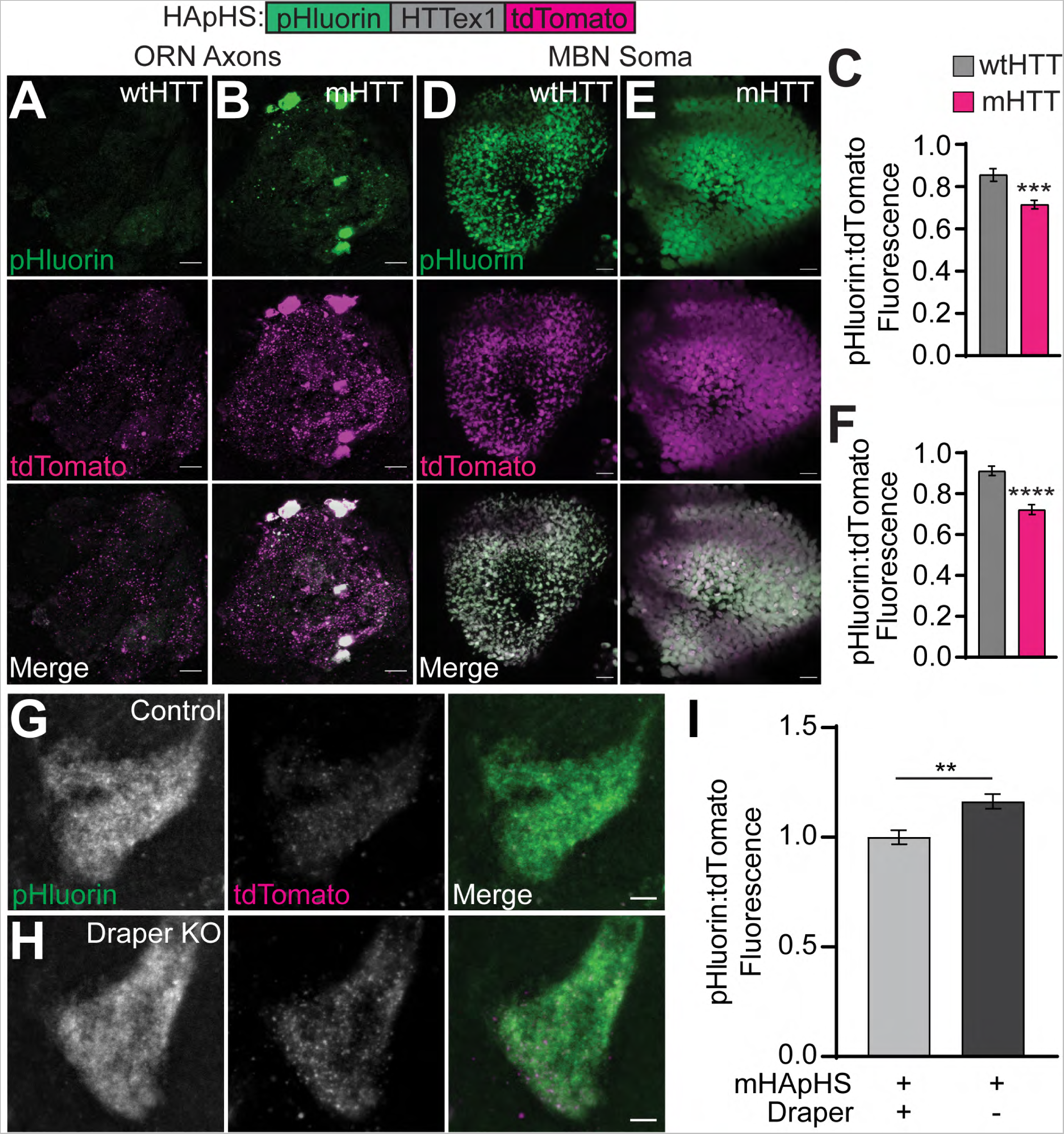
Neuronal mHTT_ex1_ accumulates in acidic cellular compartments. **(A-B, D-E)** Maximum intensity projections of (**A-B**) Or83b+ ORN axons or (**D-E**) OK107+ MBN soma expressing (**A,D**) HTT_ex1_Q25- or (**B,E**) HTT_ex1_Q91-associated pH sensor (HApHS) from 13-14 day-old flies. Scale bars = 10 μm. (**C,F**) pHluorin:tdTomato fluorescence intensity ratios of data shown in (**A-B** and **D-E**), normalized to wtHTT_ex1_ controls. (**G-H**) Maximum intensity projections of VA1lm ORN axons co-expressing mHApHS and (**G**) repo-Cas9 or (**H**) repo-Cas9 and gRNAs targeting *draper*. Scale bars = 5 μm. (**I**) pHluorin:tdTomato fluorescence intensity ratios calculated in VA1lm glomeruli from 9-10 day-old flies as shown in (**G-H**). Data were normalized to flies not expressing gRNAs. All quantified data are shown as mean ± s.e.m.; \*\**p*<0.01, \*\*\**p*<0.001, ****p<0.0001 by unpaired two-tailed *t*-test.

### Neuronal mHTT impairs injury-responsive gene upregulation

Glia alter their transcriptional profile to elicit cellular responses to insult or injury in the brain. For example, acute CNS injury in *Drosophila* increases transcription of many genes involved in phagocytosis and innate immunity, including the cell surface receptors, Draper and Toll-6, and components of their downstream signaling pathways (Fig. 5A) (Purice et al., 2017; Byrns et al., 2021; Alphen et al., 2022). To test whether the reduced ability of glia to clear mHTT_ex1_-containing axonal debris correlates with reduced injury responsiveness at a transcriptional level, we used qPCR and GFP-tagged reporters to quantify changes in gene expression of key components of these phagocytic and innate immunity pathways following mHTT_ex1_ accumulation in neurons. mHTT_ex1_ expression in uninjured Or83b+ ORNs increased relative expression of *toll-6, relish*, and *drpr-I* transcripts between 1.2- and 1.5-fold (Fig. 5B) and levels of GFP-tagged Toll-6 and Jra proteins in the CNS (Fig. 5C-H), suggesting that neuronal mHTT_ex1_ aggregates activate a mild injury response in the brain. Jra-GFP expression increased throughout the CNS and in Repo+ nuclei (Fig. 5F-H), suggesting that mHTT_ex1_-induced upregulation of this subunit of the AP-1 transcription factor can be at least partially attributed to glia. To test whether mHTT_ex1_ impairs the ability of glia to respond to acute neural injury, we monitored gene expression changes after bilateral antennal and maxillary palp nerve ablation. Similar to previous reports (Purice et al., 2017), expression of *toll-6, dorsal, relish, drpr-I*, *mmp1,* and *ets21c* genes was significantly increased 3 hrs after injury to ORN axons expressing either mCD8-GFP or wtHTT_ex1_ (Fig. 5B). However, injury-induced upregulation of each of these genes was significantly reduced in animals expressing mHTT_ex1_ in Or83b+ ORNs (Fig. 5B), suggesting that mHTT_ex1_ aggregation attenuates glial transcriptional responses to injury. We further analyzed the impact of mHTT_ex1_ on downstream immune responses in the brain by measuring induction of antimicrobial peptide (AMP) genes, well-established transcriptional targets of activated Relish/NFkB following Toll-6 or immune-deficiency pathway activation (Swanson et al., 2020b, 2020a; Alphen et al., 2022). Levels of five AMP genes, including *drosomycin*, *attacinA*, *attacinD*, *diptericinA*, and *metchnikowin*, were significantly increased 3 hours after ORN axotomy (Fig. 6A-E), similar to previous reports (Katzenberger et al., 2013; Swanson et al., 2020b; Marischuk et al., 2021). Interestingly, mHTT_ex1_ expression in ORNs alone was sufficient to induce upregulation of *drosomycin* and *attacinD* (Fig. 6A & C), albeit to a lesser extent than following acute injury. Further, injury-induced upregulation of *drosomycin* and *attacinA* was significantly reduced by mHTT_ex1_ expression in ORNs, suggesting that the activity of Relish-dependent signaling is altered by accumulation of mHTT_ex1_ aggregates. Together, these data indicate that neuronal mHTT_ex1_ aggregates trigger a mild immune response in the brain, but also inhibit the ability of glia to mount robust transcriptional responses to neural injury.

**Figure 5.**
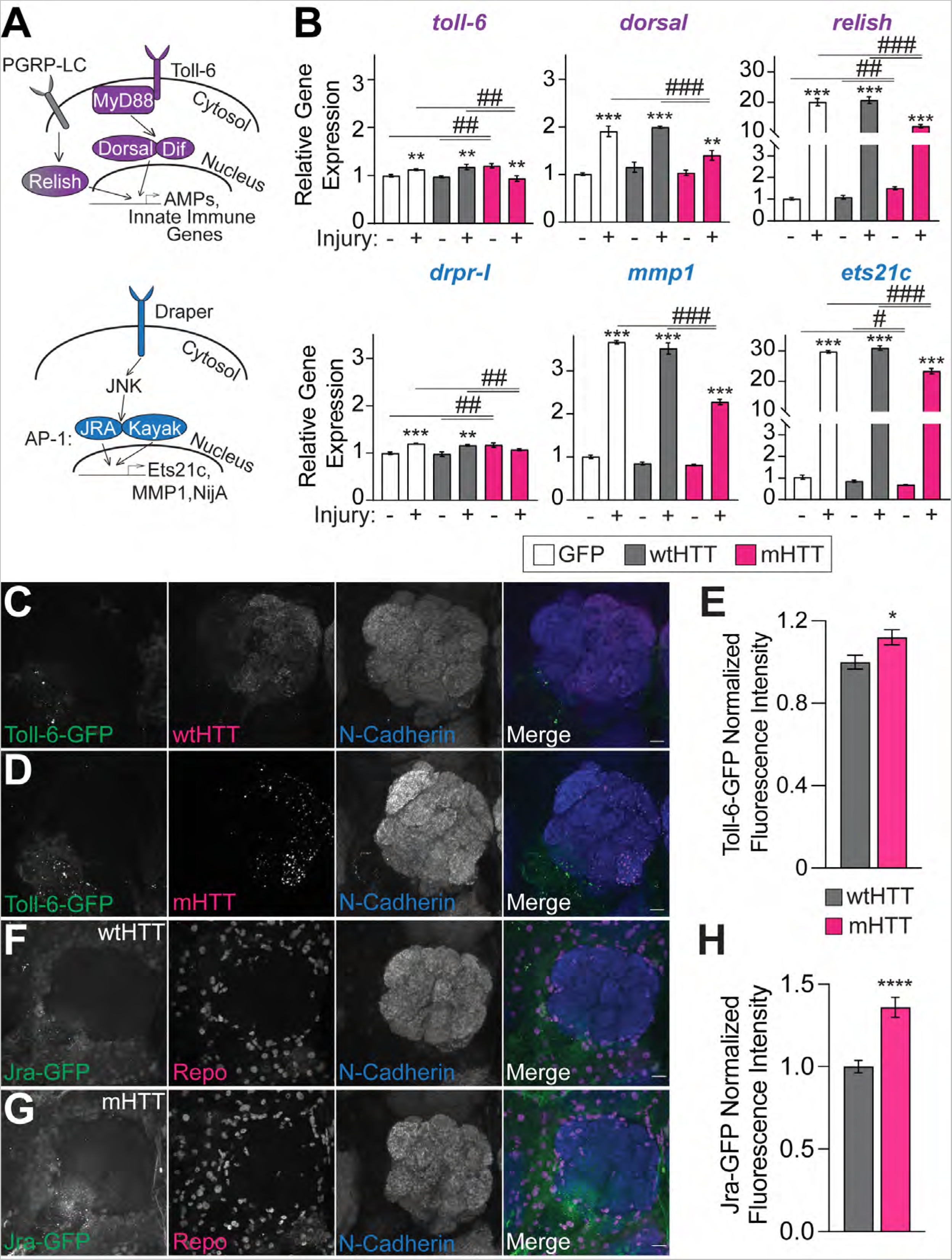
mHTT_ex1_ expression in ORNs upregulates phagocytic and innate immunity genes and impairs injury-induced transcriptional responses. (**A**) Diagrams of Toll-6 (purple) and Draper (blue) signaling pathways. (**B**) qPCR analysis of the indicated genes in 8-11 day-old flies expressing GFP, HTT_ex1_Q25-, or HTT_ex1_Q91-GFP in Or83b+ ORNs. RNA was isolated from heads of uninjured flies or flies 3 hours after bilateral antennal and maxillary palp nerve injury. Data are shown as mean ± s.e.m and normalized to the housekeeping gene *rpl32*. \**p*<0.05, \*\**p*<0.01, \*\*\**p*<0.001 by one-way ANOVA; asterisks and hashtags indicate statistical significance comparing-/+ injury or genotypes, respectively. (**C-D and F-G**) Maximum intensity projections of Or83b+ ORN axons from 14-15 day-old flies expressing (**C,F**) HTT_ex1_Q25- or (**D,G**) HTT_ex1_Q91-V5 in (**C-D**) Toll-6^MIMICGFP^ (Toll-6-GFP) or (**F-G**) Jra-GFP genetic backgrounds. Brains were immunostained with GFP, V5, and N-Cadherin (**C-D**) or GFP, Repo, and N-Cadherin (**F-G**) antibodies. In panel (**C**), diffuse wtHTT_ex1_ signal was adjusted post-acquisition for increased visibility. Scale bars = 10 μm. (**E,H**) Quantification of (**E**) Toll-6-GFP or (**H**) Jra-GFP expression from flies show in (**C-D** and **F-G**). Data are graphed as mean ± s.e.m.; \**p*<0.05, ****p<0.0001 by unpaired two-tailed t-test.

**Figure 6.**
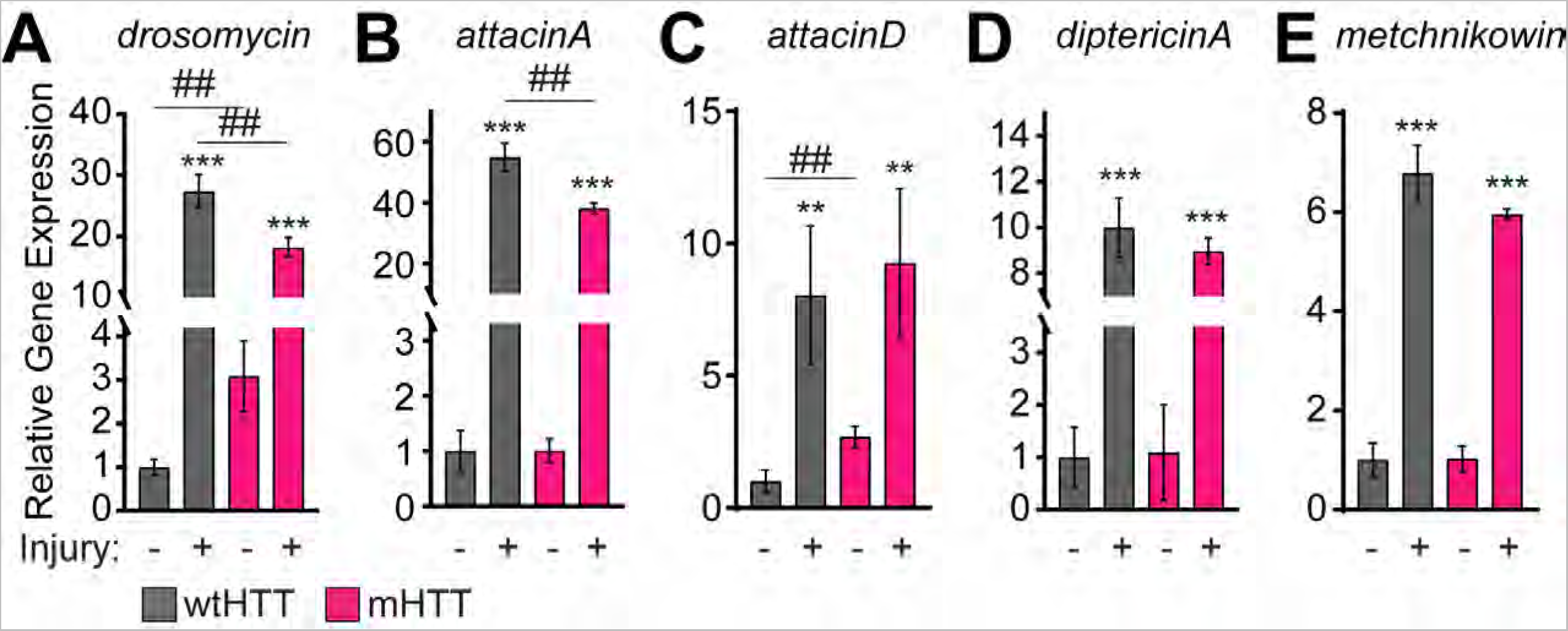
mHTT_ex1_ expression in ORNs increases expression and impairs injury-induced upregulation of some AMP genes. (**A-E**) qPCR analysis of the indicated AMP genes in 8-11 day-old flies expressing HTT_ex1_Q25- or HTT_ex1_Q91-GFP in Or83b+ ORNs. RNA was isolated from heads of uninjured flies or flies 3 hours after bilateral antennal and maxillary palp nerve injury. Data are shown as mean ± s.e.m and normalized to the housekeeping gene *rpl32*. \*\**p*<0.01, \*\*\**p*<0.001 by unpaired two-tailed t-test; asterisks and hashtags indicate statistical significance comparing-/+ injury or genotypes, respectively.

Our findings are in agreement with previously published work demonstrating that expression of mHTT_ex1_ and Aβ in neurons activates Draper-dependent phagocytosis, likely in an effort to reduce levels of these pathogenic proteins in the brain (Pearce et al., 2015; Ray et al., 2017). We further investigated this by immunostaining adult fly brains expressing mHTT_ex1_ in ORNs to monitor expression levels and localization of endogenous Draper protein. In 1 week-old adult flies expressing neuronal mHTT_ex1_, Draper immunolabeling increased ∼1.2-fold in the vicinity of ORN axons compared with age-matched controls expressing wtHTT_ex1_ (Fig. 7A-C). Closer analysis revealed that in some cases, Draper immunofluorescence was directly adjacent to or surrounding mHTT_ex1_-mCherry fluorescence (Fig. 7D). To further examine these interactions, we used image segmentation and three-dimensional reconstruction of confocal stacks to represent mHTT_ex1_-mCherry+ aggregates and Draper+ glial membranes as individual “surfaces” (Fig. 7E-F), as previously described (Donnelly et al., 2020). Close physical association of mHTT_ex1_ with Draper and other glial proteins was defined as surfaces located ≤0.2 μm from a mHTT_ex1_ object. Interestingly, whereas almost no Draper signal was detected near wtHTT_ex1_ (Fig. 7E), ∼14% of mHTT_ex1_ aggregates were closely associated with Draper surfaces (Fig. 7F1-3). These findings are consistent with Draper+ glial membranes being recruited to neuronal mHTT_ex1_ aggregates to facilitate engulfment.

**Figure 7.**
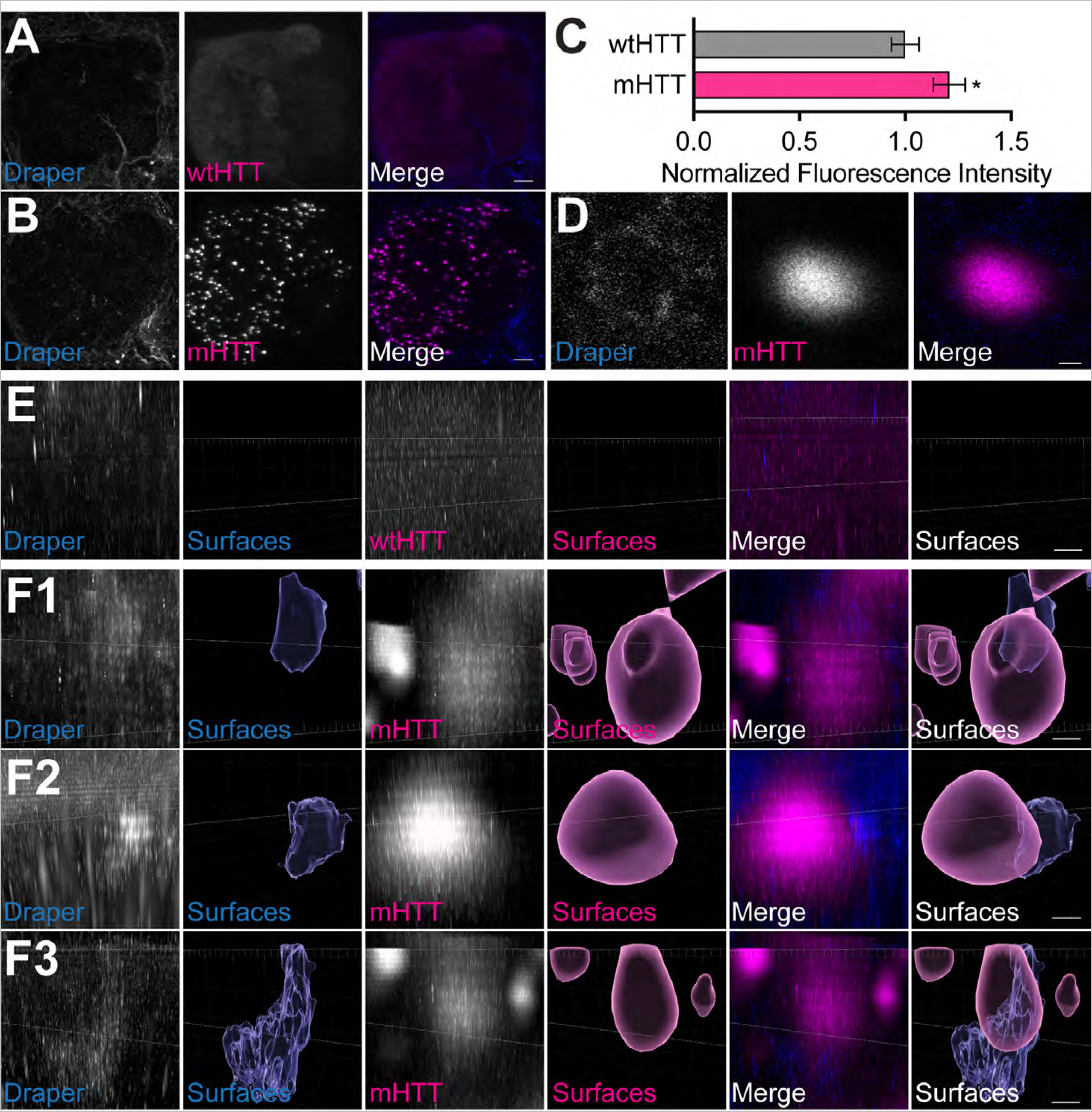
A subset of mHTT_ex1_ aggregates are closely associated with Draper+ glial membranes. (**A-B**) Maximum intensity projections of Or83b+ ORN axons from 7-8 day-old flies expressing (**A**) HTT_ex1_Q25- or (**B**) HTT_ex1_Q91-mCherry and immunostained for Draper. Scale bars = 10 μm. (**C**) Quantification of Draper immunofluorescence from flies show in (**A-B**). Data are normalized to control and graphed as mean ± s.e.m.; \**p*<0.05 by unpaired two-tailed *t*-test; n=12. (**D**) Single 0.35 μm confocal slice showing magnified HTT_ex1_Q91-mCherry aggregate and associated Draper signal. Scale bar = 0.5 μm. (**E-F**) High-magnification confocal stacks of Draper signal within ≤0.2 μm of either (**E**) HTT_ex1_Q25- or (**F1-3**) HTT_ex1_Q91-mCherry surfaces. Raw data are shown to the left of segmented surfaces generated from each fluorescence signal. In (**E**), diffuse wtHTT_ex1_ signal was adjusted post-acquisition for increased visibility. Scale bars = 1 μm.

Synaptic neuropil in the *Drosophila* brain is primarily inhabited by two glial subtypes, ensheathing glia and astrocytic glia (Doherty et al., 2009). In adult flies, ensheathing glia compartmentalize synaptic regions and respond to CNS injury by upregulating Draper and clearing debris via phagocytosis (Doherty et al., 2009). Consistent with our previous findings in all repo+ glia (Pearce et al., 2015), mz0709+ ensheathing glia were vulnerable to prion-like conversion of glial wtHTT_ex1_ proteins by mHTT_ex1_ aggregates generated in DA1 ORN axons (Fig. 8A, C-D). Seeded wtHTT_ex1_ aggregates were defined as wtHTT_ex1_ signal that colocalized with mHTT_ex1_ objects identified by image segmentation of confocal stacks (Donnelly et al., 2020). Conversely, seeding of wtHTT_ex1_ expressed in alrm+ astrocytic glia, which lack detectable Draper expression in the adult fly brain (Doherty et al., 2009), was not observed (Fig. 8B-D). Interestingly, ensheathing glial-specific RNAi knockdown of Toll-6, Relish, and NijA increased mHTT_ex1_ aggregate numbers in DA1 ORN axons, similar to the effects of Draper-I knockdown (Fig. 9A-B). Further, adult-specific, pan-glial knockdown of Ets21c, which was found to be required for normal development, also increased numbers of mHTT_ex1_ aggregates in DA1 ORN axons (Fig. 9C-D). Mean mHTT_ex1_ aggregate volume was not affected by ensheathing glial knockdown of these genes except for a ∼20% increase following NijA depletion (Fig. 9E), suggesting that aggregate or vesicle size was unaffected by these genetic manipulations. Thus, several glial genes with established roles in phagocytic and innate immune signaling regulate basal turnover of mHTT_ex1_ aggregates in ORN axons.

**Figure 8.**
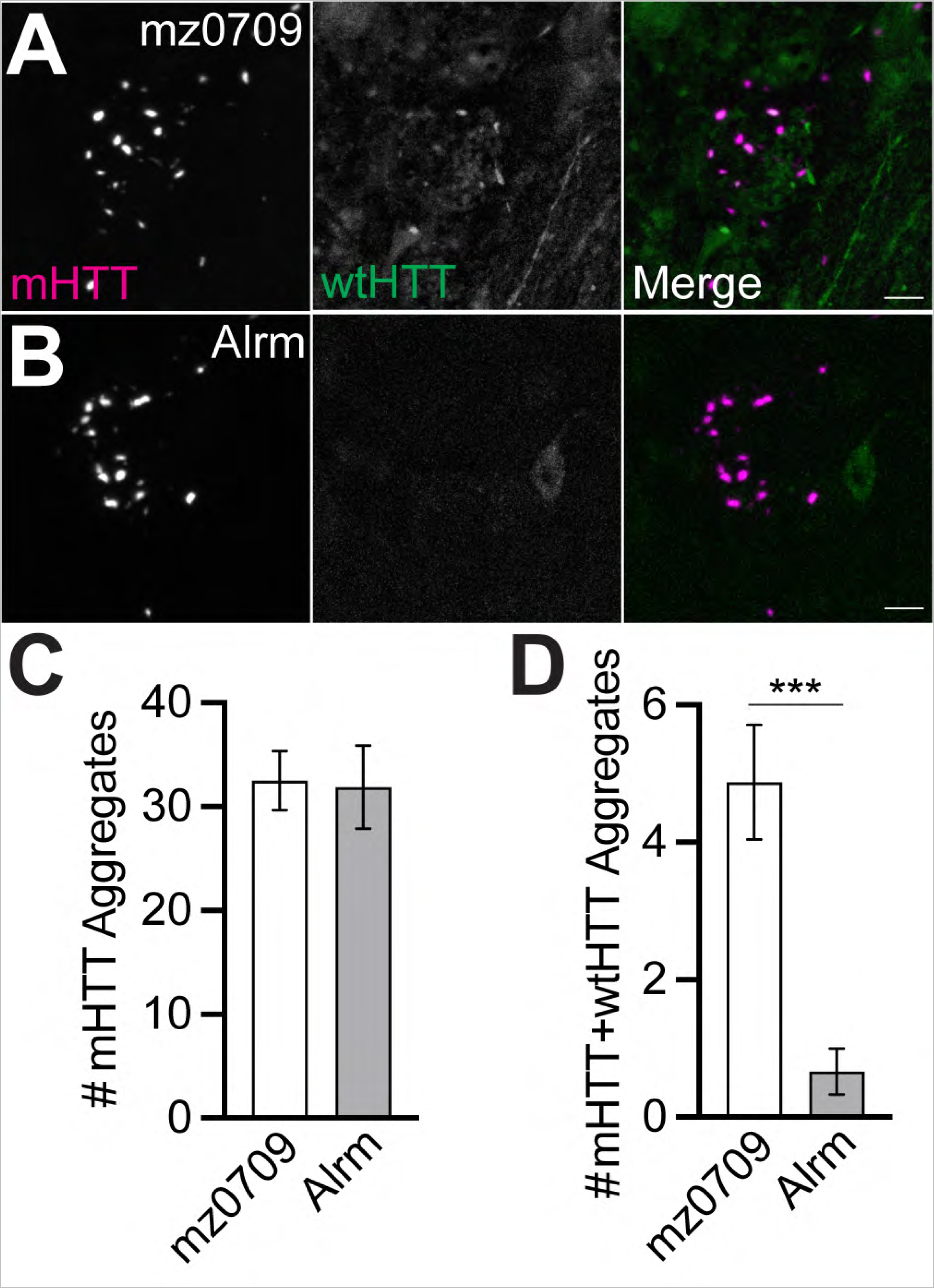
Seeded aggregation of wtHTT_ex1_ protein expressed in ensheathing glia by neuronal mHTT_ex1_ aggregates. (**A-B**) Maximum intensity projections of DA1 glomeruli from 4-5 day-old flies expressing HTT_ex1_Q91-mCherry in DA1 ORNs and HTT_ex1_Q25-GFP in (**A**) mz0709+ ensheathing glia or (**B**) Alrm+ astrocytic glia. Scale bars = 5 μm. (**C-D**) Quantification of (**C**) HTT_ex1_Q91-mCherry (“mHTT”) and (**D**) seeded HTT_ex1_Q25-GFP (“mHTT+wtHTT”) aggregates from flies shown in (**A-B**). Data are shown as mean ± s.e.m.; \*\*\**p*<0.001 by unpaired two-tailed *t*-test.

**Figure 9.**
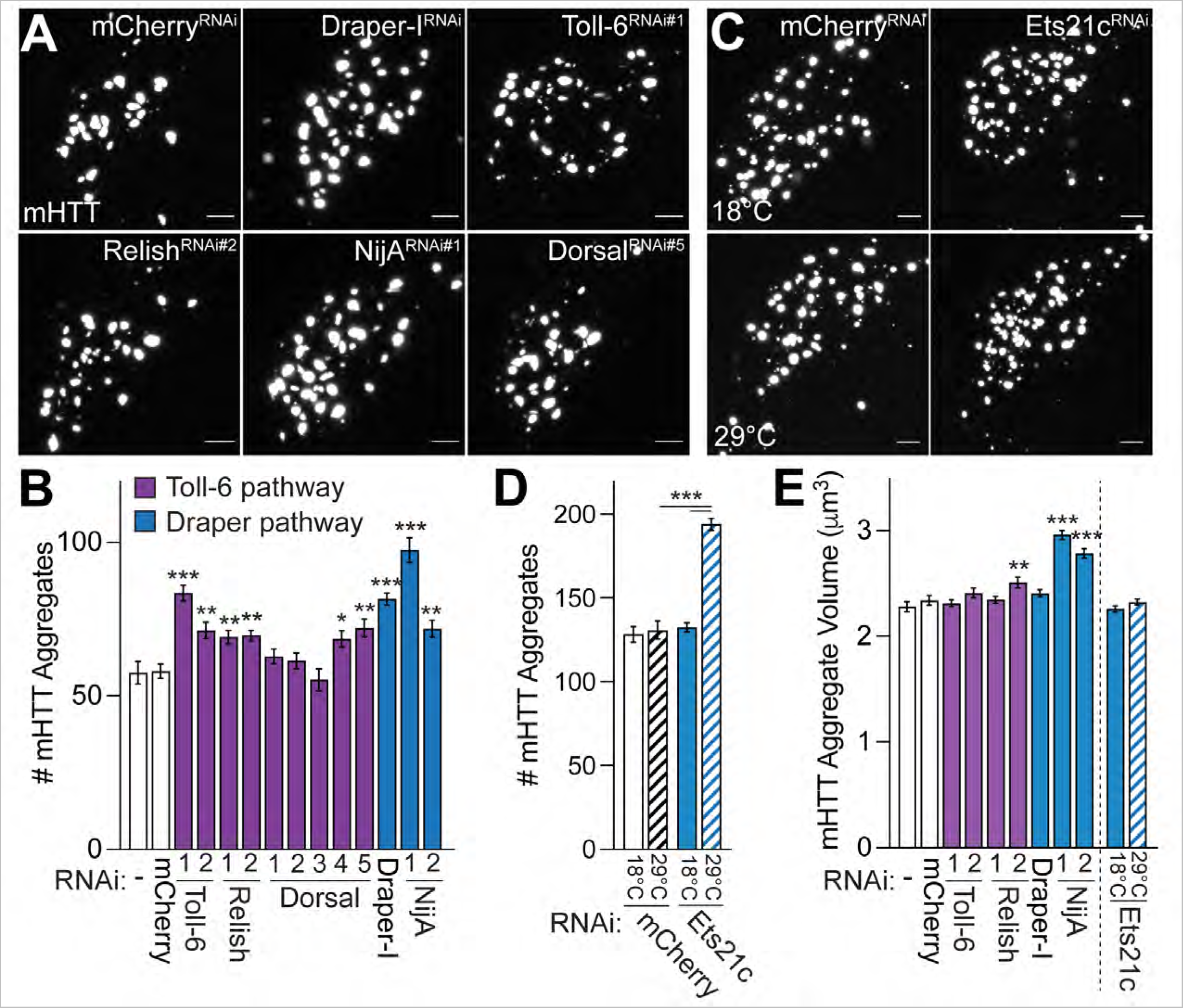
Glial phagocytic and innate immunity genes regulate numbers of mHTT_ex1_ aggregates in ORN axons. (**A**) Maximum intensity projections of DA1 glomeruli from 7 day-old flies expressing HTT_ex1_Q91-mCherry in DA1 ORNs and siRNAs targeting the indicated genes in ensheathing glia. Scale bars = 5 μm. (**B)** Quantification of HTT_ex1_Q91-mCherry aggregates detected in DA1 glomeruli from flies shown in (**A**). (**C**) Maximum intensity projections of DA1 glomeruli from 7 day-old flies expressing HTT_ex1_Q91-mCherry in DA1 ORNs and siRNAs targeting mCherry or Ets21c in repo+ glia in the presence of tubP-Gal80^ts^. Adult flies were raised at the permissive (18°C, *top*) or restrictive (29°C, *bottom*) temperatures to regulate siRNA expression in adults. Scale bars = 5 μm. (**D**) Quantification of HTT_ex1_Q91-mCherry aggregates detected in DA1 glomeruli from flies shown in (**C**). (**E**) Mean volumes for HTT_ex1_Q91-mCherry aggregates detected in DA1 glomeruli from flies shown in (**A and C**). All graphed data are shown as mean ± s.e.m.; \**p*<0.05, \*\**p*<0.01, \*\*\**p*<0.001 by one-way ANOVA or unpaired two-tail *t*-test compared to no RNAi or mCherry^RNAi^ controls.

### Neuronal mHTT aggregates are associated with defects in multiple endolysosomal compartments

Our findings thus far indicate that neuronal mHTT_ex1_ aggregates elicit mild injury responses in the brain and reduce the ability of phagocytic glia to respond transcriptionally and functionally to acute nerve injury. We have previously postulated that prion-like spreading of mHTT_ex1_ in the fly CNS could be facilitated by escape of engulfed aggregates from the endolysosomal compartment of phagocytic glia (Pearce et al., 2015; Donnelly et al., 2020). Therefore, we sought to determine whether neuronal mHTT_ex1_ aggregates are associated with defects in endolysosomal processing in non-injured brains. We first measured effects of neuronal mHTT_ex1_ expression on the quantity, size, and function of lysosomes using dyes that label active cathepsins and low pH cellular compartments. Expression of mHTT_ex1_ in Or83b+ ORNs increased numbers of lysosomes labeled by the active cathepsin dye, Magic Red (MR) (Fig. 10A-C), and the low pH sensor, LysoTracker Red (LTR) (Fig. 10D-F), compared with control brains expressing wtHTT_ex1_. High resolution analysis of confocal stacks and filtering for segmented MR+ and LTR+ surfaces within 0.2 μm of a mHTT_ex1_ object revealed close association of MR+ and LTR+ signals with mHTT_ex1_ aggregates (Fig. 10C and F-H).

**Figure 10.**
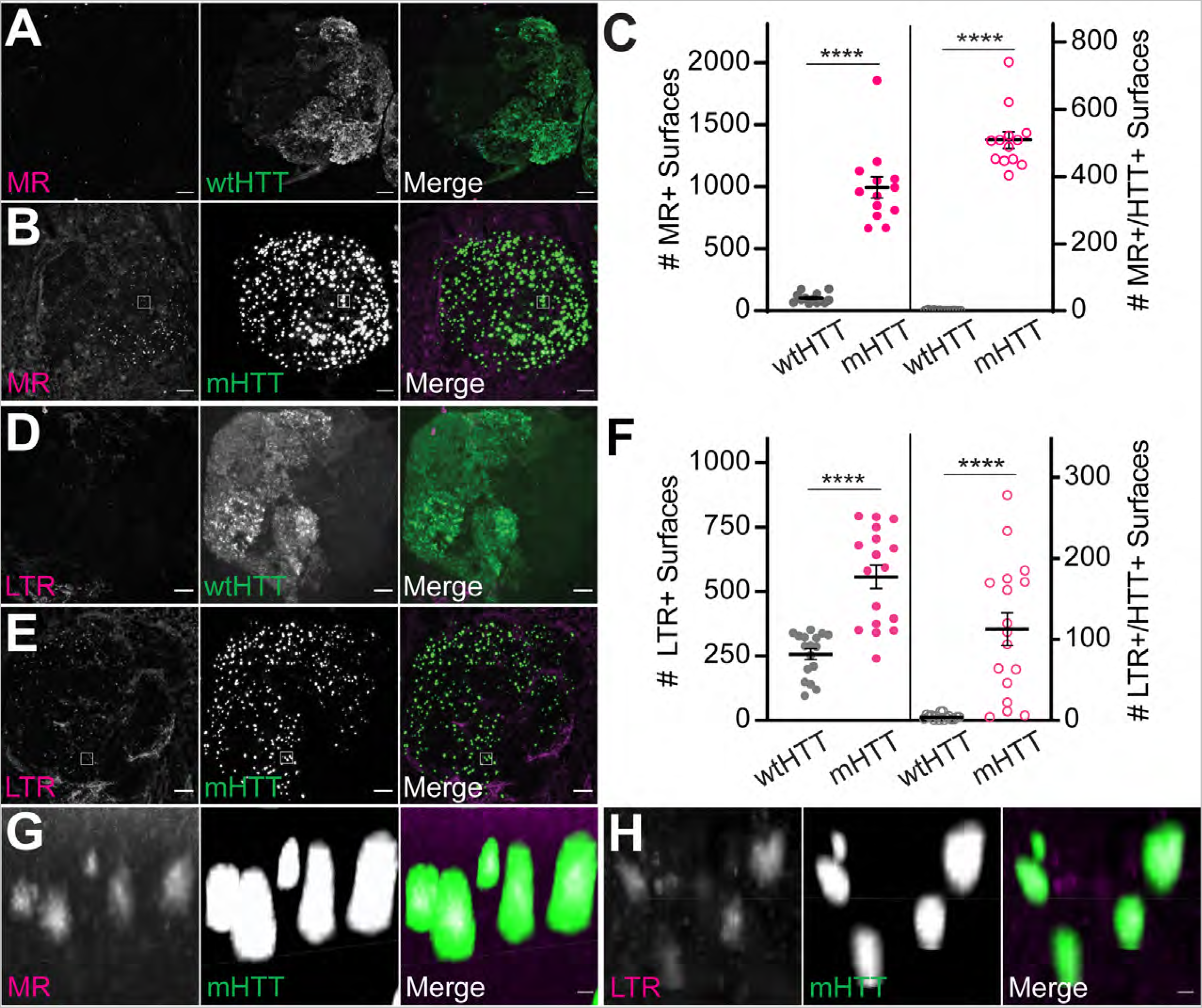
Neuronal mHTT_ex1_ expression increases numbers of acidified and active lysosomes in adult brains. (**A-B**) Maximum intensity projections of antennal lobes from 9-10 day-old adult flies expressing (**A**) HTT_ex1_Q25- or (**B**) HTT_ex1_Q91-GFP in Or83b+ ORNs and stained with Magic Red (MR) to label active cathepsins. Scale bars = 10 μm. (**C**) Quantification of MR+ surfaces (*left*) and HTT_ex1_-associated MR+ surfaces (*right*) from flies shown in (**A** and **B**); HTT_ex1_-associated vesicles were defined by filtering for MR+ surfaces ≤0.2 μm from HTT_ex1_ fluorescent signal in confocal stacks. **(D-E**) Maximum intensity projections of antennal lobes from 15 day-old flies expressing (**D**) HTT_ex1_Q25- or (**E**) HTT_ex1_Q91-GFP in Or83b+ ORNs and stained with Lysotracker Red (LTR) to label low pH compartments. Scale bars = 10 μm. Diffuse wtHTT_ex1_ signal was adjusted post-acquisition for increased visibility in panels (**A** and **D**). (**F**) Quantification of LTR+ surfaces (*left*) and HTT_ex1_-associated LTR+surfaces (*right*) from flies shown in (**D** and **E**); HTT_ex1_-associated vesicles were defined by filtering for LTR+ surfaces ≤0.2 μm from HTT_ex1_ fluorescent signal in confocal stacks. Data are shown as mean ± s.e.m.; \*\*\*\**p*<0.0001 by unpaired two-tailed *t*-test. (**G-H**) High-magnification images of boxes indicated in (**B and E**) showing co-localization of MR (**G**) or LTR (**H**) with mHTT_ex1_ fluorescent signals. Scale bars = 1 μm.

We next employed *Drosophila* genetic tools to assess the impact of neuronal mHTT_ex1_ specifically on glial lysosomes by driving expression of lysosomal-associated membrane protein 1 (LAMP1) tagged at its cytosolic C-terminus with GFP (LAMP1-GFP) in all glia. Neuronal mHTT_ex1_ expression increased the overall number of glial LAMP1-GFP+ vesicles and the number of LAMP1-GFP+ vesicles in close proximity to HTT_ex1_ signal (Fig. 11A-C and G-H). We typically observed only partial overlap of LAMP1-GFP signal with mHTT_ex1_ aggregate surfaces identified by this method, possibly due to incomplete labeling or rupture of lysosomal membranes as a result of aggregate size or structural features. Interestingly, this subpopulation of LAMP1-GFP+ lysosomes closely associated with mHTT_ex1_ were enlarged compared with all lysosomes (Fig. 11I). To examine whether neuronal mHTT_ex1_ aggregates affect lysosome integrity, we expressed a transgene encoding LAMP1 fused at its N-terminus to GFP in glia. This construct integrates into the lysosomal membrane such that GFP is exposed to the lumen, and loss of GFP signal can thus be used to monitor LAMP1+ lysosome degradative activity (Pulipparacharuvil et al., 2005). Interestingly, the quantity and mean volume of GFP-LAMP1+ vesicles were significantly increased in brains expressing mHTT_ex1_ in Or83b+ ORNs (Fig. 11D-G), suggesting lysosomal enlargement and dysfunction due to mHTT_ex1_ aggregates. GFP-LAMP1+ surfaces were also more associated with mHTT_ex1_ aggregates compared to wtHTT_ex1_ controls (Fig. 11F and I).

**Figure 11.**
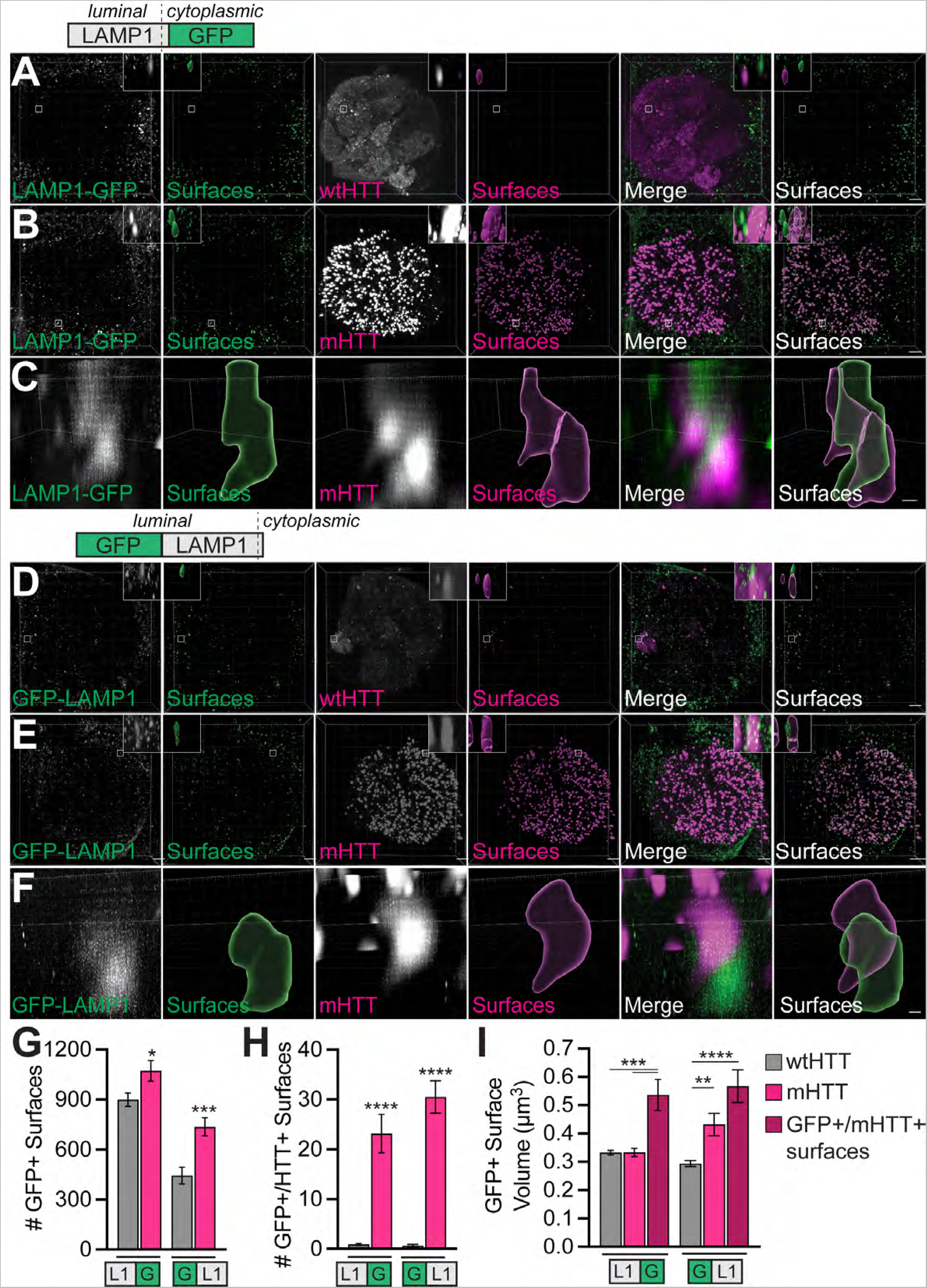
LAMP1+ vesicle accumulation in fly brains expressing mHTT_ex1_ in ORNs. (**A-B**) Confocal stacks showing antennal lobes from 19-22 day-old flies expressing (**A**) HTT_ex1_Q25- or (**B**) HTT_ex1_Q91-mCherry in Or83b+ ORNs and LAMP1 tagged at its cytoplasmic C-terminus with GFP (LAMP1-GFP) in glia. Brains were immunostained with anti-GFP to amplify LAMP1-GFP signal. LAMP1-GFP+ or HTT_ex1_+ segmented surfaces are shown to the right of each set of raw fluorescence images. Insets show magnified regions of interest from each image. Scale bars = 10 μm. (**C**) High-magnification confocal stack showing a LAMP1-GFP+ surface within 0.2 μm of two HTT_ex1_Q91-mCherry+ aggregates. Scale bar = 1 μm. **(D-E**) Confocal stacks showing antennal lobes from 21-22 day-old flies expressing (**D**) HTT_ex1_Q25- or (**E**) HTT_ex1_Q91-mCherry in Or83b+ ORNs and LAMP1 tagged at its luminal N-terminus with GFP (GFP-LAMP1) in glia. GFP-LAMP1+ or HTT_ex1_+ segmented surfaces are shown to the right of each set of raw fluorescence images. Insets show a single GFP-LAMP1+ surface of interest from each image. Scale bars = 10 μm. Diffuse wtHTT_ex1_ signal was adjusted post-acquisition for increased visibility in panels (**A** and **D**). (**F**) High-magnification confocal stack showing a GFP-LAMP1+ surface within 0.2 μm of a HTT_ex1_Q91-mCherry+ aggregate. Scale bar = 1μm. (**G-I**) Quantification of total LAMP1-GFP+ or GFP-LAMP1+ surfaces (**G**), LAMP1-GFP+ or GFP-LAMP1+ surfaces ≤0.2 μm from HTT_ex1_ surfaces (**H**), and mean LAMP1-GFP+ or GFP-LAMP1+ surface volume (**I**) in brains expressing HTT_ex1_Q25- or HTT_ex1_Q91-mCherry. The dark red bars in (**I**) represent LAMP1+ surfaces that co-localized with mHTT_ex1_. All graphed data are shown as mean ± s.e.m.; \**p*<0.05, \*\**p*<0.01, \*\*\**p*<0.001, \*\*\*\**p*<0.0001 by unpaired two-tailed *t*-test.

To directly test whether glial endolysosomes experience increased membrane damage due to neuronal mHTT_ex1_ expression, we generated transgenic flies expressing mCherry-tagged Galectin-3 or Galectin-8, lectins that translocate from the cytoplasm to the lumen of ruptured lysosomes and endosomes, respectively (Aits et al., 2015; Daussy and Wodrich, 2020; Jia et al., 2020). Neuronal mHTT_ex1_ expression increased overall numbers of glial Galectin-3/8+ surfaces (Fig. 12A-E), Galectin-3/8+ surfaces closely associated with mHTT_ex1_ aggregates (Fig. 12F), and mean volume of mHTT_ex1_-associated Galectin-3/8+ vesicles (Fig. 12G). Together, these data suggest that neuronal mHTT_ex1_ aggregates induce non cell-autonomous accumulation, enlargement, and membrane damage of endolysosome vesicles in glial cells.

**Figure 12.**
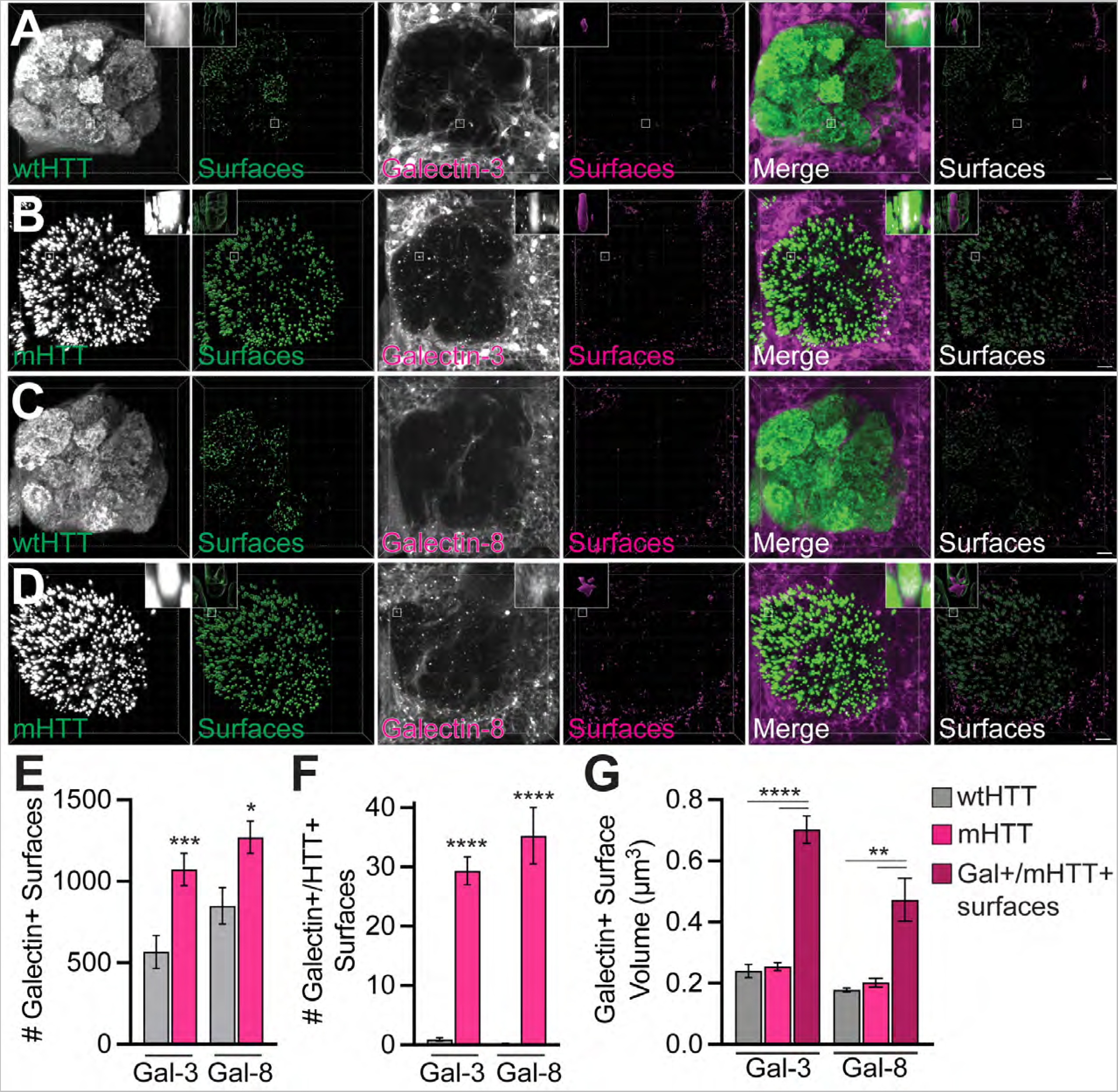
Increased association of Galectins-3 and -8 expressed in glia with neuronal mHTT_ex1_ aggregates. (**A-D**) Confocal stacks showing antennal lobes from 16-18 day-old flies expressing HTT_ex1_Q25-(**A and C**) or HTT_ex1_Q91-GFP (**B and D**) in Or83b+ ORNs together with Galectin-3 (**A-B**) or Galectin-8 (**C-D**) tagged with mCherry in glia. Segmented Galectin+ or HTT_ex1_+ surfaces are shown to the right of each set of raw fluorescence images. Insets show Galectin+ surfaces of interest from each image. Scale bars = 10 μm. (**E-G**) Quantification of total Galectin+ surfaces (**E**), Galectin+ surfaces ≤0.2 μm from HTT_ex1_ surfaces (**F**), and mean Galectin+ surface volume (**G**) in brains expressing HTT_ex1_Q25- or HTT_ex1_Q91-mCherry. The dark red bars in (**G**) represent Galectin+ surfaces that co-localized with mHTT_ex1_. All graphed data are shown as mean ± s.e.m.; \**p*<0.05, \*\**p*<0.01, \*\*\**p*<0.001, \*\*\*\**p*<0.0001 by unpaired two-tailed *t*-test.

“Seeding-competent” mHTT aggregates are defined by their ability to nucleate or “seed” the aggregation of normally-soluble wtHTT proteins, a defining characteristic of infectious prion and other prion-like proteins (Jucker and Walker, 2018; Donnelly et al., 2022). Many studies have pointed to a role for defective clearance by endolysosomal pathways in promoting the propagation of pathogenic aggregates (Freeman et al., 2013; Jiang et al., 2017; Chen et al., 2019; Jiang and Bhaskar, 2020; Polanco and Götz, 2022). To test whether altered glial lysosome function affects seeding competency of mHTT_ex1_, we used RNAi to individually knockdown proteins with known roles in lysosome degradation in glia and examined the ability of neuronal mCherry-tagged mHTT_ex1_ to seed aggregation of glial, GFP-tagged wtHTT_ex1_. Depletion of two subunits of the vacuolar ATPase (V-ATPase), Vha68-3 (Portela et al., 2018) and rabconnectin-3A (Yan et al., 2009), and Spinster, a late-endosomal and lysosomal efflux permease (Rong et al., 2011), increased numbers of glial wtHTT_ex1_ aggregates detected as GFP+ surfaces that colocalized with mHTT_ex1_ aggregates (Fig. 13A-C) (Donnelly et al., 2020). Knockdown of Vha16-1, a V-ATPase subunit that regulates endolysosome membrane fusion (Finbow et al., 1994; Dunlop et al., 1995), did not affect wtHTT_ex1_ seeding; however, it did cause accumulation of mHTT_ex1_ aggregates in DA1 ORN axons (Fig. 13A and B). Together, these findings suggest that disruption of normal glial lysosome acidification and/or degradative capacity promotes formation of seeding-competent mHTT_ex1_ aggregates.

**Figure 13.**
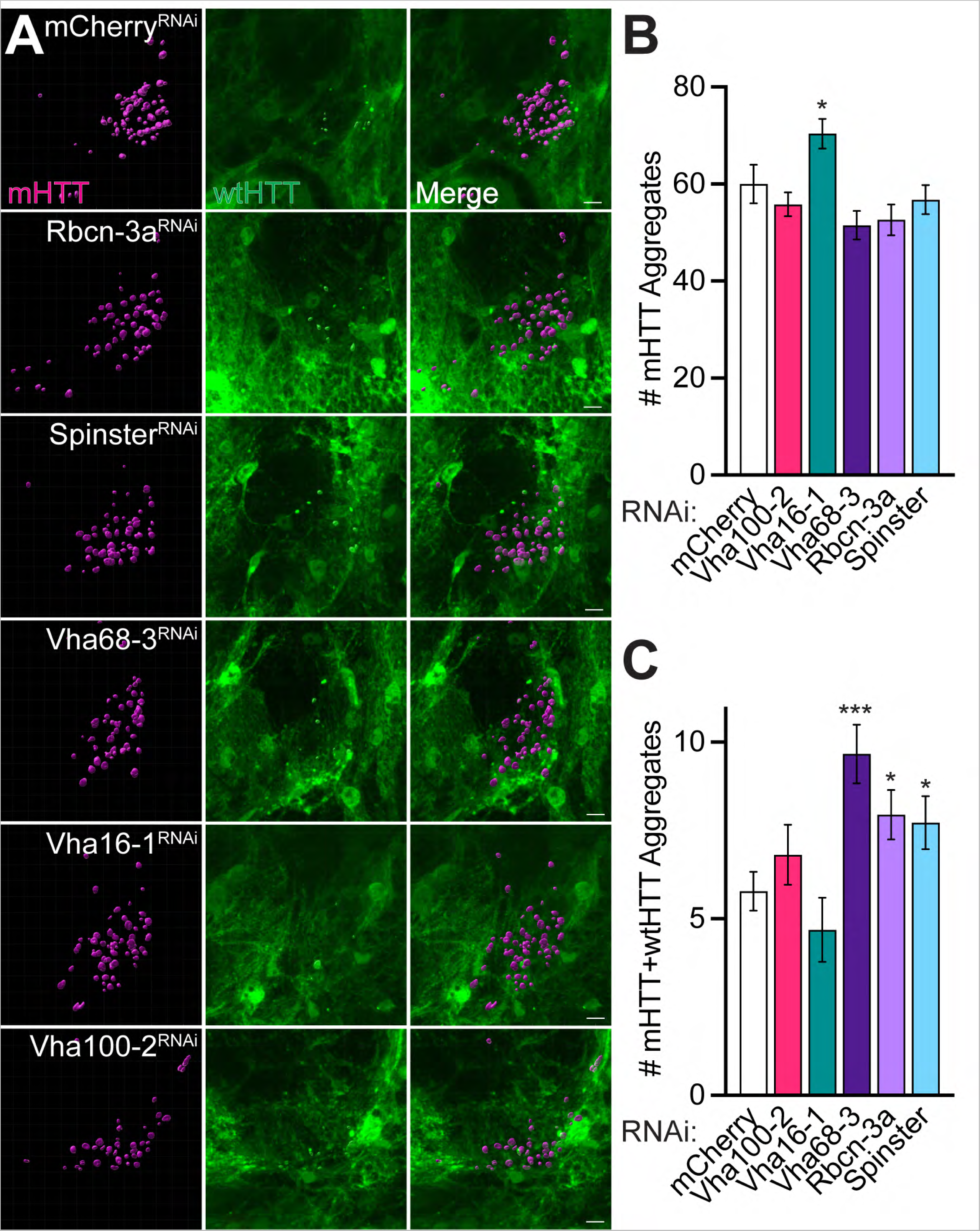
Knockdown of genes regulating lysosome acidification alters seeded aggregation of glial wtHTT_ex1_ protein by neuronal mHTT_ex1_ aggregates. (**A**) Confocal stacks showing DA1 glomeruli from 8-9 day-old flies expressing HTT_ex1_Q91-mCherry in DA1 ORNs and HTT_ex1_Q25-GFP plus siRNAs targeting the indicated genes in repo+ glia. Negative controls expressed siRNAs targeting mCherry. Scale bars = 5 μm. (**B-C**) Quantification of (**B**) HTT_ex1_Q91-mCherry or (**C**) seeded HTT_ex1_Q25-GFP aggregates from brains shown in (**A**). Data are shown as mean ± s.e.m.; \**p*<0.05, \*\*\**p*<0.005 by unpaired two-tailed *t*-test.

### The GTPase Rab10 mediates prion-like transmission of mHTT aggregates

Lysosome dysfunction could occur secondary to upstream defects in endo/phagosome maturation. Our prior work supports a model in which a portion of mHTT_ex1_ aggregates engulfed by glia evade degradation during phagosome maturation and/or phagolysosome formation (Pearce et al., 2015; Donnelly et al., 2020). To test this model, we used forward genetic screening to interrogate roles for glial Rab GTPases in prion-like conversion of cytoplasmic wtHTT_ex1_ proteins by engulfed neuronal mHTT_ex1_ aggregates. The *Drosophila* genome encodes 31 Rab and Rab-like proteins, all of which have mammalian orthologs, and most of these GTPases are implicated in vesicle and target membrane fusion in cells (Zhang et al., 2007). To determine whether any *Drosophila* Rabs mediate escape of phagocytosed mHTT_ex1_ aggregates and seeding of wtHTT_ex1_ in the glial cytoplasm, we individually knocked down each Rab in repo+ glia using RNAi. Glial-restricted silencing of 23 of the 31 Rabs produced viable adults, and these flies were used to monitor effects of Rab knockdown on mHTT_ex1_-induced aggregation of wtHTT_ex1_ in glia. Only two Rab RNAi lines, *Rab10^RNAi^*and *Rab23^RNAi#2^*, significantly altered numbers of induced wtHTT_ex1_ aggregates (Fig. 14A-C). Of note, 3 additional Rab23 RNAi lines had no significant effects on numbers of wtHTT_ex1_ aggregates, suggesting that *Rab23^RNAi#2^* may cause off-target effects. Strikingly, Rab10 depletion reduced numbers of seeded wtHTT_ex1_ aggregates, phenocopying effects of Draper knockdown (Fig. 14A and C) and suggesting that Draper and Rab10 function in the same pathway. To test whether this effect of Rab10 loss-of-function was mediated via a reduction in Draper expression, we measured endogenous Draper immunofluorescence in *rab10* null flies. Draper protein levels were ∼18% lower in *rab10 ^-/-^* animals compared to wild-type controls (Fig. 14D). This reduction is unlikely to fully account for decreased seeding of wtHTT_ex1_ following Rab10 knockdown, as numbers of wtHTT_ex1_ aggregates are similar between wild-type and *draper* heterozygotes (Fig. 14G) (Pearce et al., 2015). To further explore this, we attempted to restore Draper function via overexpression of Draper-I, which rescues loss of Draper function in aged flies (Purice et al., 2016). However, transgenic expression of Draper-I in glia failed to rescue the effects of Rab10 knockdown on wtHTT_ex1_ aggregate seeding (Fig. 14E). These findings suggest that Rab10 acts downstream of Draper rather than by regulating Draper activity. We further tested for interactions between *drpr* and *rab10* using loss-of-function alleles to examine effects on mHTT_ex1_ aggregate transmission from presynaptic DA1 ORNs to postsynaptic projection neurons (PNs), a process previously reported to require transport though Draper+ glia (Donnelly et al., 2020). Because reduced survival of *rab10* null flies (Kohrs et al., 2021) was exacerbated by transgenic HTT_ex1_ expression, we tested for genetic interaction between *drpr* and *rab10* in heterozygous and trans-heterozygous animals. While we detected no change in mHTT_ex1_ aggregate numbers (Fig. 14F), significantly fewer wtHTT_ex1_ aggregates formed in PNs of *rab10^+/-^ drpr^+/-^* transheterozygotes than in individual heterozygotes (Fig. 14G). This same effect was not observed in *rab14^+/-^ drpr^+/-^* transheterozygous animals (Fig. 14G), indicating that the genetic interaction is specific to *drpr* and *rab10*. Thus, Draper and Rab10 appear to function in the same phagocytic pathway that regulates prion-like transmission of phagocytosed mHTT_ex1_ aggregates in the fly brain.

**Figure 14.**
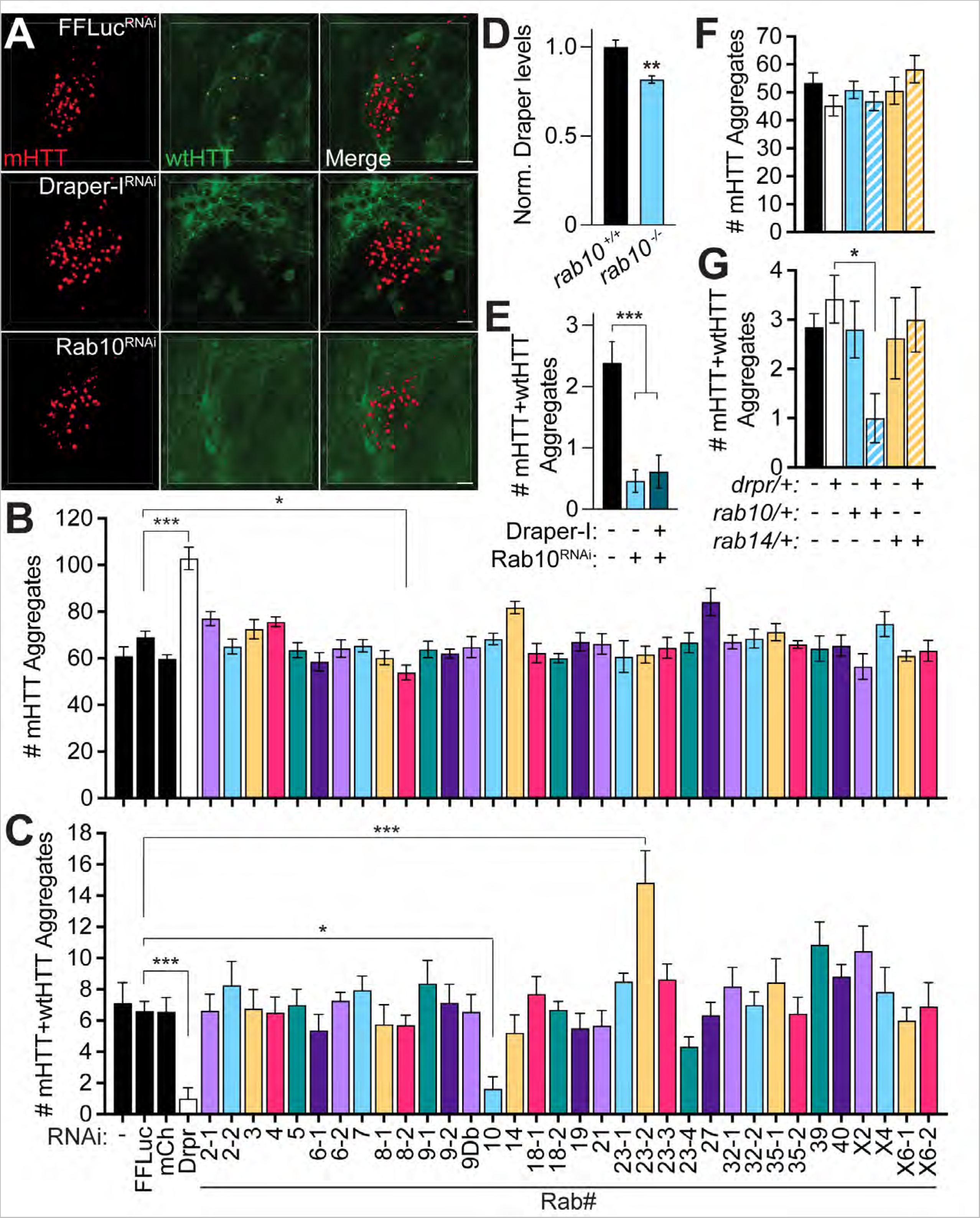
Rab10 is required for seeded aggregation of glial wtHTT_ex1_ by neuronal mHTT_ex1_ aggregates. (**A**) Confocal stacks of DA1 glomeruli from 7 day-old flies expressing HTT_ex1_Q91-mCherry in DA1 ORNs and HTT_ex1_Q25-GFP plus siRNAs targeting firefly luciferase (FFLuc), Draper, or Rab10 in repo+ glia. Surfaces representing mHTT_ex1_ aggregates (*red*) and seeded wtHTT_ex1_ aggregates are superimposed on the raw data. Scale bars = 5μm. (**B-C**) mHTT_ex1_ (**B**) or mHTT_ex1_+wtHTT_ex1_ (**C**) aggregates quantified from flies expressing HTT_ex1_Q91-mCherry in DA1 ORNs and HTT_ex1_Q25-GFP plus siRNAs targeting Draper-I (Drpr) or 23 different Rab GTPases in repo+ glia. Negative controls expressed no siRNAs or siRNAs targeting FFLuc or mCherry (*black bars*). (**D**) Normalized Draper immunofluorescence in the central brain of 4-5 day-old wild-type (*rab10 ^+/+^*) and *rab10* null (*rab10 ^-/-^*) flies. (**E**) Quantification of seeded wtHTT_ex1_ aggregates in flies expressing HTT_ex1_Q91-mCherry in DA1 ORNs and HTT_ex1_Q25-GFP plus siRNAs targeting Rab10, without or with Draper-I cDNAs in repo+glia. (**F-G**) Quantification of mHTT_ex1_ (**F**) or mHTT_ex1_+wtHTT_ex1_ (**G**) aggregates from 10 day-old flies expressing HTT_ex1_Q91-mCherry in DA1 ORNs and HTT_ex1_Q25-GFP in PNs, either heterozygous or trans-heterozygous for *draper* (*drpr/+*), *rab10* (*rab10/+*), or *rab14* (*rab14/+*) mutant alleles. All graphed data are shown as mean ± s.e.m.; \**p*<0.05, \*\**p*<0.01, \*\*\**p*<0.005 by one-way ANOVA or unpaired two-tailed *t-*test comparing to controls.

### Neuronal mHTT aggregates alter numbers of early and late glial phagosomes

Our forward genetic screen indicates that at least one glial Rab GTPase, Rab10, regulates the seeding capacity of neuronal mHTT_ex1_ aggregates. Interestingly, Rab10 has been reported to regulate phagosome maturation in mammalian cells (Cardoso et al., 2010; Seto et al., 2011; Lee et al., 2020; Wang et al., 2023), and Rab10 expression and activity are altered in neurodegenerative diseases, including AD and PD (Eguchi et al., 2018; Tavana et al., 2018; Yan et al., 2018). Because little is known about Rab10’s role in glia, we sought to characterize this GTPase alongside two additional Rabs with well-established roles in endocytosis, Rab5 and Rab7, markers of early and late endo/phagosomes, respectively (Hutagalung and Novick, 2011). Interestingly, we found that *rab10*, *rab5*, and *rab7* were upregulated between ∼1.4 and 2.4-fold following acute injury to ORN axons, and mHTT_ex1_ expression in Or83b+ ORNs alone caused upregulation of *rab10* and *rab7* genes by ∼1.2-fold (Fig. 15A).

**Figure 15.**
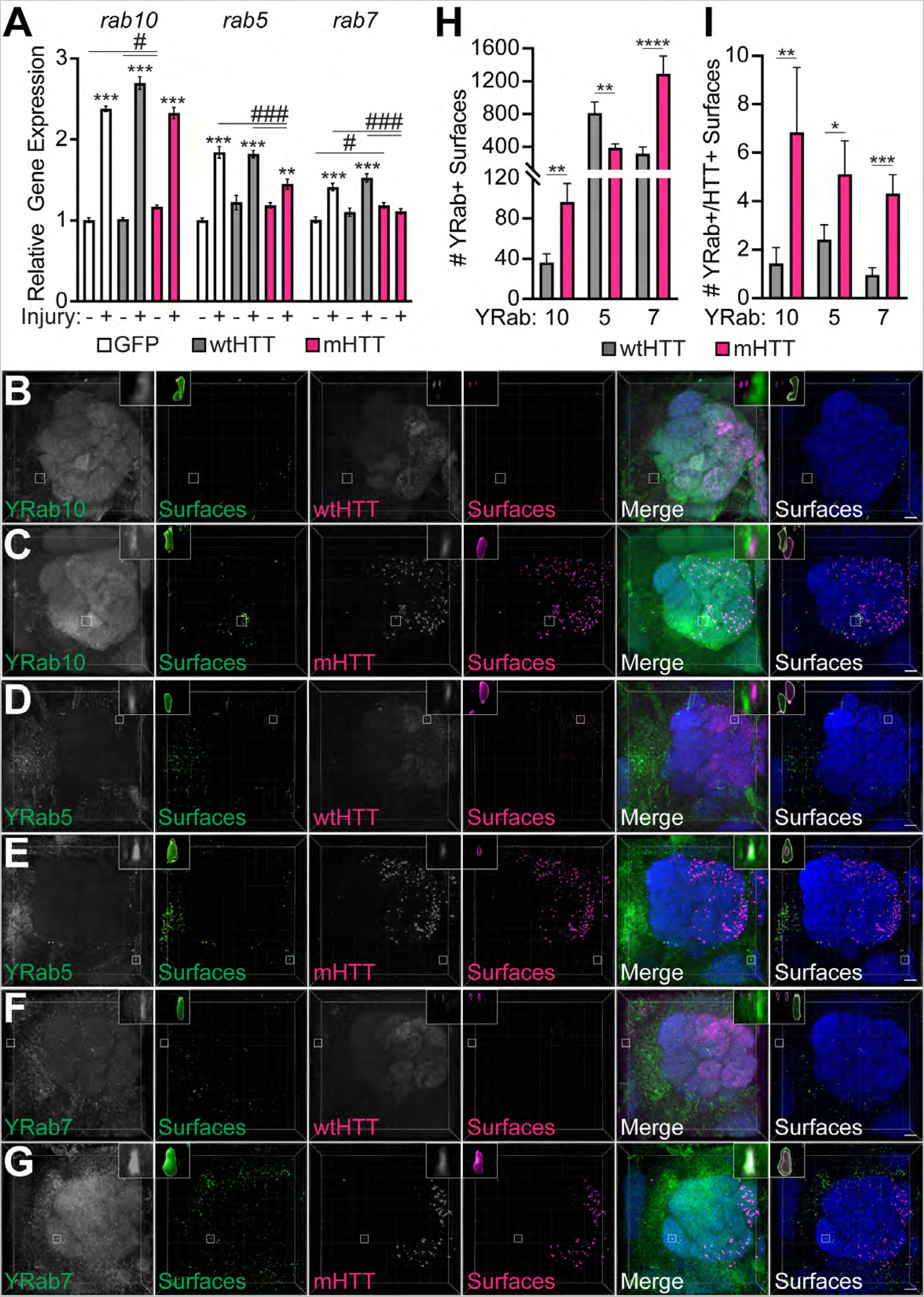
Association of neuronal mHTT_ex1_ aggregates with Rab GTPases that label early, maturing, and late phagosomes. (**A**) qPCR analysis of *rab10*, *rab5*, and *rab7* expression in 8-11 day-old flies expressing GFP, HTT_ex1_Q25-, or HTT_ex1_Q91-GFP in Or83b+ ORNs. RNA was isolated from heads of uninjured flies or flies 3 hours after bilateral antennal and maxillary palp nerve injury. Data are shown as mean ± s.e.m and normalized to the housekeeping gene *rpl32*. \**p*<0.05, \*\**p*<0.01, \*\*\**p*<0.001 by one-way ANOVA, asterisks and hashtags indicate statistical significance comparing-/+ injury or genotypes, respectively. (**B-G**) Confocal stacks of the antennal lobe from 7 day-old flies expressing (**B, D, & F**) HTT_ex1_Q25- or (**C, E, & G**) HTT_ex1_Q91-V5 in Or83b+ ORNs and endogenously-tagged (**B-C**) YRab10, (**D-E**) YRab5, or (**F-G**) YRab7 in all cells. Brains were immunostained using YFP (*green*), V5 (*magenta*), and N-Cadherin (*blue*) antibodies. Segmented YRab+ or HTT+ surfaces are shown to the right of each set of raw fluorescent images. Insets show magnified YFP+ surfaces of interest from each image. Diffuse wtHTT_ex1_ signal was adjusted post-acquisition for increased visibility in panels (**B, D,** and **F**). Scale bars = 10 μm. (**H-I**) Quantification of YRab+ surfaces (**H**) and YRab+ surfaces within 0.2μm of HTT_ex1_+ surfaces (**I**). Data are shown as mean ± s.e.m.; \**p*<0.05, \*\**p*<0.01, \*\*\**p*<0.001, \*\*\*\**p*<0.0001 by unpaired two-tailed *t*-test.

Interestingly, injury-induced upregulation of *rab5* and *rab7* was inhibited by ∼50% and ∼100%, respectively, in flies expressing mHTT_ex1_ compared to controls (Fig. 15A). Altogether, these results identify *rab5*, *rab7*, and *rab10* as novel injury-response genes in the fly CNS and suggest that neuronal mHTT_ex1_ aggregates impair injury-induced responses of *rab5* and *rab7*.

We next examined effects of neuronal mHTT_ex1_ expression on the localization of Rab10, Rab5, and Rab7 proteins endogenously-tagged with YFP-myc at their N-termini, herein referred to as YRab10, YRab5, and YRab7 (Dunst et al., 2015). YRab+ vesicles were identified in confocal stacks as segmented YFP+ surfaces with a mean diameter of 0.3-8 μm (Fig. 15B-G), consistent with vesicle sizes reported in other fly tissues (Prince et al., 2019). Expression of mHTT_ex1_ in Or83b+ ORN axons caused an overall increase in numbers of YRab10+ and YRab7+ vesicles, but a decrease in the number of YRab5+ vesicles compared with wtHTT_ex1_ controls (Fig. 15H). Further, each of these YRab+ vesicles subpopulations were closely associated with mHTT_ex1_ aggregates more frequently than with wtHTT_ex1_ (Fig. 15I), suggesting that mHTT_ex1_ protein interacts with each of these intracellular vesicle subpopulations in the brain.

To assess effects of neuronal mHTT_ex1_ aggregates specifically on glial Rab+ compartments, we expressed YFP-tagged Rab5, 7, and 10 transgenes in all glia. Similar to our findings with endogenous YRabs, expression of mHTT_ex1_ in Or83b+ ORN axons was associated with increased numbers of glial YFP-Rab10+ and -Rab7+ vesicles (Fig. 16A-B, E-F, and G); however, we observed a significant decrease in glial YFP-Rab5+ vesicle abundance (Fig. 16C-D and G). YFP-Rab10+, -5+, and -7+ vesicles increased their association with axonal mHTT_ex1_ aggregates compared with wtHTT_ex1_ (Fig. 16H), in many cases with closely associated YFP-Rab+ signal partially surrounding a mHTT_ex1_ aggregate (Fig. 17). Of note, only a small fraction of YFP-Rab+ vesicles were identified as associated with mHTT_ex1_ aggregates, possibly due to transient interactions with vesicle compartments as aggregates transit the glial phagolysosomal system, heterogeneous labeling of phagosomes by these markers in intact brain tissue, or because our selection filter excluded YFP-Rab+ surfaces located >0.2 μm away from an aggregate. The mean volume of YFP-Rab7+ vesicles that interacted with mHTT_ex1_ aggregates was significantly increased compared with wtHTT_ex1_ controls (Fig. 16I), suggesting that mHTT_ex1_ leads to enlargement of Rab7+ late phagolysosomes. Together, these data suggest that accumulation of phagocytosed mHTT_ex1_ aggregates in Rab7+ or Rab10+ late phagosomes and decreased association with early Rab5+ phagosomes could be a key mechanism underlying protein aggregate-induced toxicity and spreading in HD.

**Figure 16.**
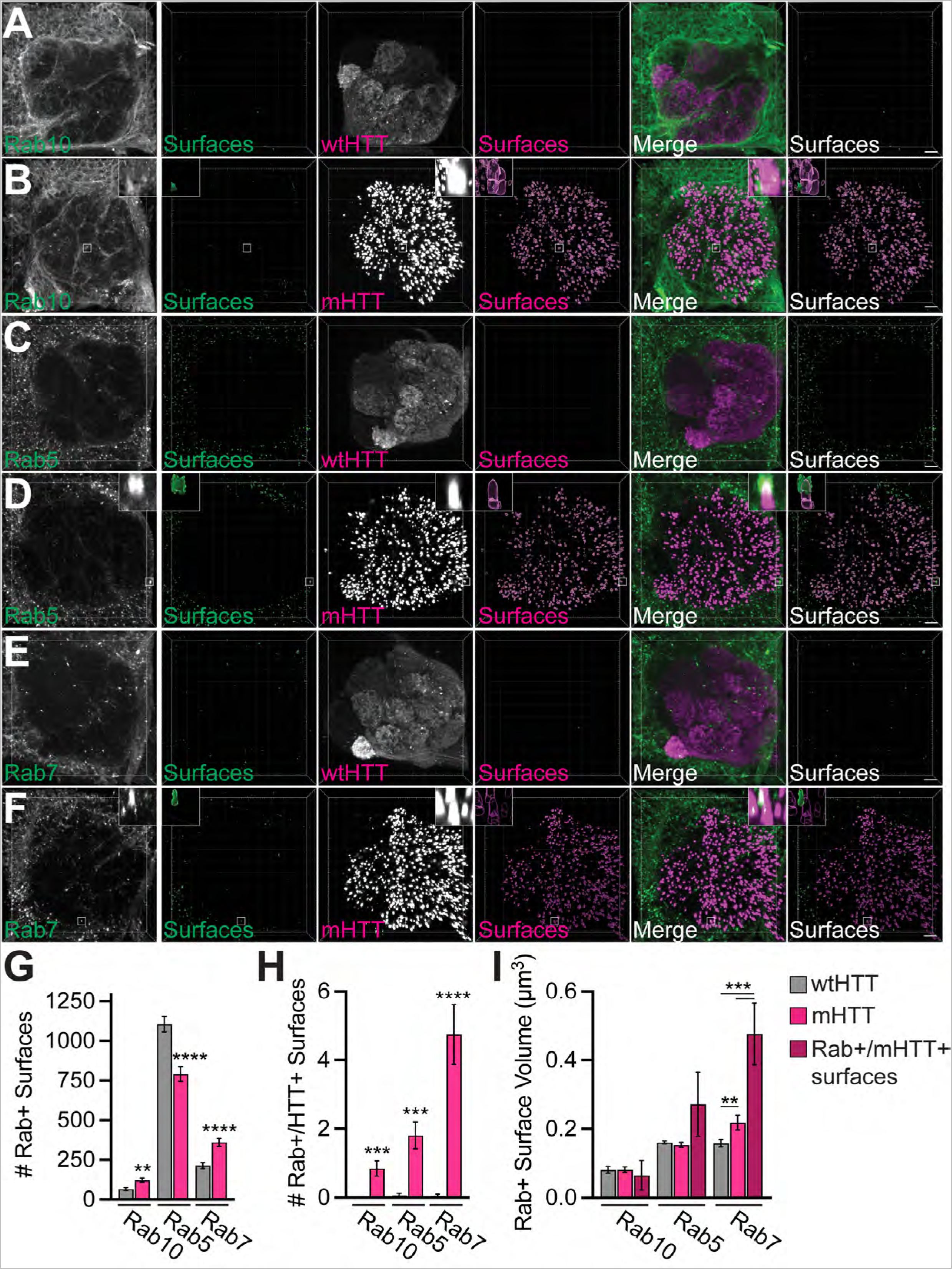
Association of neuronal mHTT aggregates with glial Rab GTPases that label early, maturing, and late phagosomes. (**A-F**) Confocal stacks of the antennal lobe from 16-19 day-old flies expressing (**A, C, E**) HTT_ex1_Q25- or (**B, D, F**) HTT_ex1_Q91-V5 in Or83b+ ORNs together with (**A-B**) YFP-Rab10, (**C-D**) YFP-Rab5, or (**E-F**) YFP-Rab7 in repo+ glia. Brains were immunostained with anti-GFP to amplify YFP-Rab signals. Segmented Rab+ or HTT_ex1_+ surfaces are shown to the right of each set of raw fluorescence images. Insets show magnified YFP-Rab+ surfaces of interest from each image. Diffuse wtHTT_ex1_ signal was adjusted post-acquisition for increased visibility in panels (**A, C,** and **E**). Scale bars = 10 μm. (**G-I**) Quantification of total YFP-Rab+ surfaces (**G**), YFP-Rab+ surfaces ≤0.2 μm from HTT_ex1_ surfaces (**H**), and mean YFP-Rab+ surface volume (**I**) in brains expressing HTT_ex1_Q25- or HTT_ex1_Q91-mCherry. The dark red bars in (**I**) represent YFP-Rab+ surfaces that co-localized with mHTT_ex1_. All graphed data are shown as mean ± s.e.m.; \**p*<0.05, \*\**p*<0.01, \*\*\**p*<0.001, \*\*\*\**p*<0.0001 by unpaired two-tailed *t*-test.

**Figure 17.**
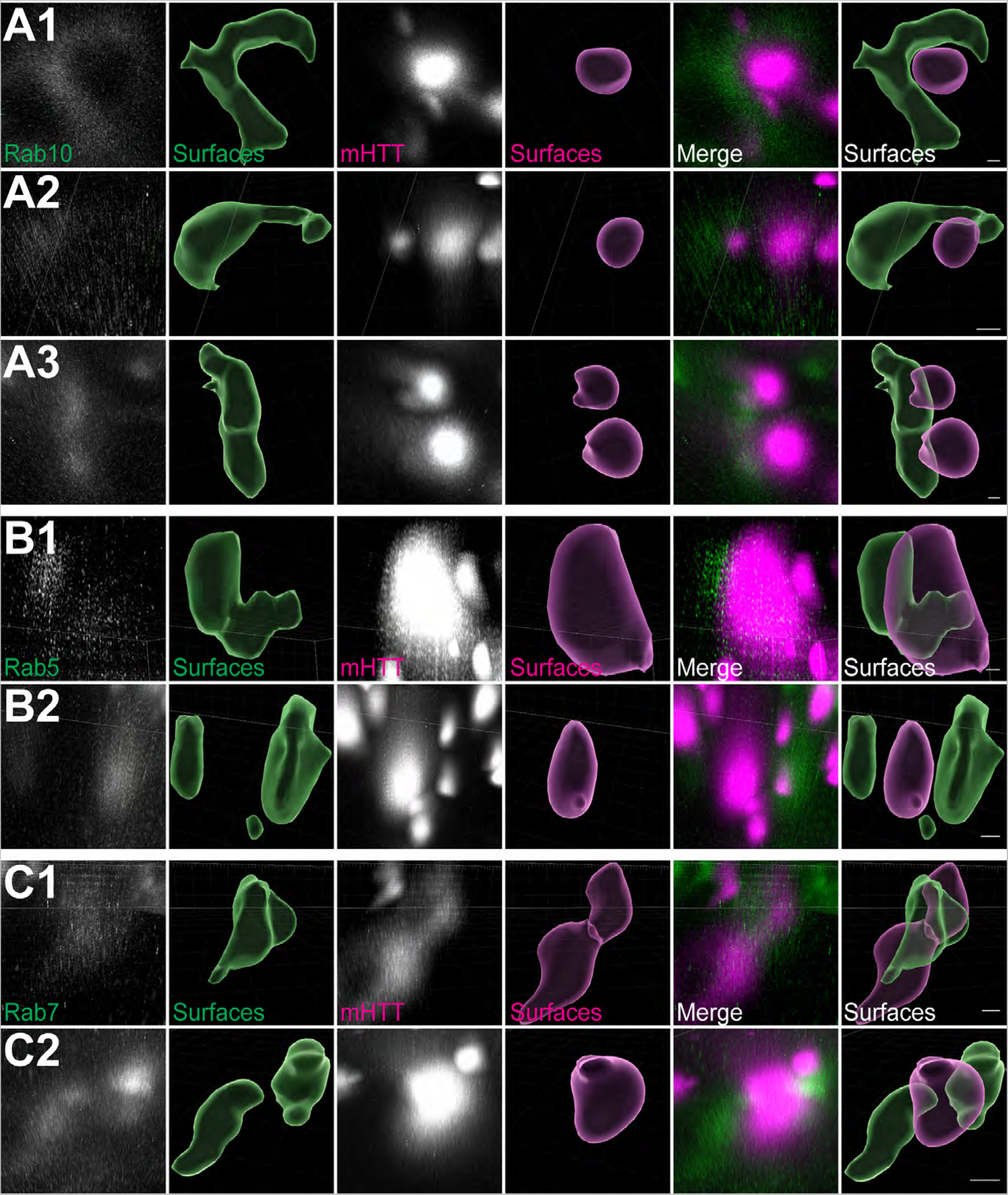
Glial YFP-Rab+ surfaces are closely associated with neuronal mHTT_ex1_ aggregates. High magnification confocal stacks showing examples of individual YFP+ surfaces within 0.2 μm of HTT_ex1_Q91 aggregates from 21-22 day-old adult brains expressing HTT_ex1_Q91-mCherry in Or83b+ ORNs and YFP-Rab10 (**A1-3**), YFP-Rab5 (**B1-2**), or YFP-Rab7 (**C1-2**) in repo+ glia. Brains were immunostained with anti-GFP to amplify YFP-Rab signals. Scale bars = 1 μm.

## DISCUSSION

Toxic amyloid aggregates have been a primary target in neurodegenerative disease drug development for decades, with some recent promise using immunotherapy to reduce aggregate loads in the brain (Karran and De Strooper, 2022). Microglia and astrocytes have also emerged as attractive therapeutic targets in efforts to boost neuroprotective glial functions or reduce neuroinflammation.

However, approaches that target glial cells must effectively strike a balance between amplifying beneficial and reducing harmful effects of these cells in the brain. Here, we tested for interactions between phagocytic glia and pathogenic protein aggregates in a *Drosophila* model of HD. We report that aggregates formed by mHTT_ex1_ protein fragments impair glial transcriptional and functional responses to CNS injury, induce upregulation of stress response and innate immunity genes, and alter numbers of endolysosomal vesicles detected in uninjured brains. A targeted forward genetic screen revealed that Rab10, a GTPase previously reported to regulate phagosome maturation, mediates prion-like conversion of cytoplasmic wtHTT_ex1_ proteins by phagocytosed mHTT_ex1_ aggregates. Together, these findings suggest that neuronal mHTT_ex1_ aggregates compromise intracellular membrane integrity as they transit endolysosomal systems, generating toxic, seeding-competent aggregates that propagate disease phenotypes.

Glia respond to neural injury by altering their transcriptional, morphological, and metabolic profiles to promote neuronal survival and clear debris from the brain; however, failure of glia to return to a resting state elicits harmful neuroinflammatory consequences (Liddelow et al., 2020). We have previously reported that activated phagocytic glia can have both beneficial (i.e., elimination of toxic aggregates) and harmful (i.e., as vectors for aggregate spread) effects in the brain (Pearce et al., 2015; Donnelly et al., 2020). We report here that key glial injury-responsive pathways, i.e., Draper-mediated phagocytosis and Toll-6-mediated innate immune signaling, are induced in the presence mHTT_ex1_ aggregation in the adult fly brain. These findings are in line with studies from other labs demonstrating that *Drosophila* Toll-6 and mammalian Toll-Like Receptor signaling pathways are upregulated in response to dying neurons during development (McLaughlin et al., 2019) and in patient and mammalian models of neurodegenerative disease (Casula et al., 2011; Miron et al., 2018; Kouli et al., 2020). Interestingly, increased microglial NF-kB signaling mediates tau spread and toxicity in mice, further linking innate immunity to prion-like mechanisms of disease progression (Wang et al., 2022). Thus, activation of glial immune pathways may contribute to feed-forward mechanisms involving aggregate formation, pathology propagation, and neuroinflammatory signaling.

Genome-wide association studies have revealed numerous genes associated with increased risk of AD and other neurodegenerative diseases, and many of these risk variants are enriched in pathways that control key glial cell functions. For example, rare risk-associated variants of the microglial *TREM2* gene alter amyloid aggregate accumulation and seeding in cells and animal models (Leyns et al., 2019; Parhizkar et al., 2019; Jain et al., 2023). A number of additional genes involved in endolysosomal processing are associated with increased risk of AD, PD, FTD, and/or ALS, such as the phagocytic receptor *CD33*, endosomal genes *BIN1* and *RIN3*, and *GRN*, which encodes the lysosomal progranulin protein (Podleśny-Drabiniok et al., 2020; Welikovitch et al., 2023). Although Draper/MEGF10 variants are not known risk factors in neurodegenerative disease, MEGF10 is highly expressed in phagocytic astrocytes (Chung et al., 2013), mediates Aý aggregate engulfment (Singh et al., 2010; Fujita et al., 2020), and acts as a receptor for C1q, a mediator of early synapse loss in AD mouse models (Hong et al., 2016; Iram et al., 2016). Our finding that upregulation of *draper* and other phagocytic genes is inhibited by mHTT_ex1_ expression suggests that glial responsiveness and phagocytic capacity is attenuated in the presence of protein aggregate pathology in neurons. Genetic or environmental risk factors that impact glial health could accelerate these defects and exacerbate aggregate-induced neurotoxicity in the brain.

Lysosomes are essential for cell survival, not only to clear damaged or toxic materials from cells, but also as intracellular centers for macromolecule recycling, energy metabolism, and cell-cell communication. Lysosomal abnormalities, including vesicle enlargement, deacidification, and membrane leakiness, have been reported in patient brains and animal and cell models of multiple neurodegenerative diseases, suggesting that these defects play a central role in disease progression (Bonam et al., 2019; Polanco and Götz, 2022; Udayar et al., 2022). We report here that Rab7+, Rab10+, and Lamp1+ phagolysosomes accumulate in glia as a result of mHTT_ex1_ expression in neurons and that lysosome dysfunction and membrane permeabilization increase in the presence of mHTT_ex1_ aggregates. We also observed that neuronal aggregates were associated with decreased nascent phagosome formation and reduced numbers of early (Rab5+) phagosomes in glia. These findings are consistent with a “traffic jam” model (Small et al., 2017) in which mHTT_ex1_ aggregates accumulate over time in glial endolysosomal compartments, preventing proper flow of materials into (i.e., engulfment) and out of (i.e., degradation) the pathway (Fig. 18). We postulate that persistence of aggregates in faulty endolysosomes promotes formation and/or release of degradation-resistant aggregates with enhanced toxicity and seeding capacity. This hypothesis is in agreement with recent studies in tauopathy models showing that Aý aggregation occurs secondary to lysosome deacidification and membrane permeabilization (Lee et al., 2022), and hypophagocytic glia contribute to tau aggregate propagation (Hopp et al., 2018; Brelstaff et al., 2021). Release of seeding-competent aggregates from dysfunctional lysosomes could occur via active exocytosis, perhaps in an effort to alleviate cell toxicity, or passively due to vesicle rupture (Flavin et al., 2017; Falcon et al., 2018; Yuste-Checa et al., 2021).

**Figure 18.**
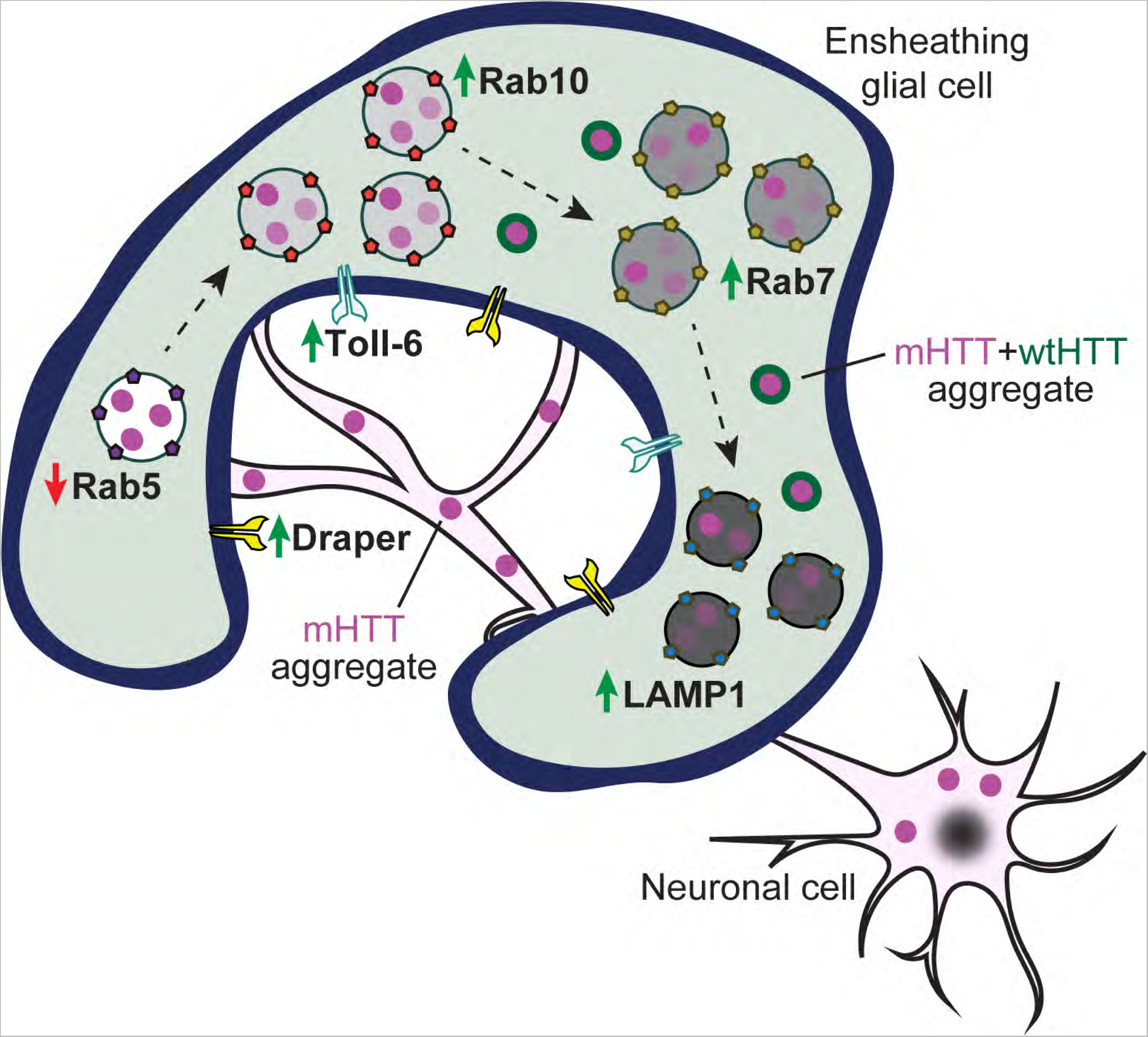
“Traffic jam” model illustrating effects of neuronal mHTT_ex1_ aggregates on phagocytic glia. mHTT_ex1_ aggregates (*magenta circles*) generated in axons cause phagolysosomal defects in neighboring glial cells. Neuronal mHTT_ex1_ aggregates activate glial Draper and Toll-6 signaling pathways, but impair normal phagocytic responses to injury, including reduced nascent phagosome formation and decreased numbers of Rab5+ early phagosomes (*red arrow*). A buildup of engulfed mHTT_ex1_ aggregates in glia leads to accumulation of maturing (Rab10+ or Rab7+) phagosomes and lysosomes (LAMP1+) (*green arrows*), possibly further slowing Draper-dependent engulfment and early phagosome formation. Defective phagocytic clearance could enhance leak or ejection of some mHTT_ex1_ aggregates from phagolysosomes to increase their toxicity and capacity to seed soluble HTT proteins (*magenta + green circles*).

Among the growing list of genes linked to neurodegenerative disease pathogenesis are Rab GTPases and genes that modify Rab functions (Kiral et al., 2018). Rab proteins are essential for vesicle sorting and trafficking in all cells and use GTP hydrolysis to differentially associate with and organize intracellular membranes (Homma et al., 2021). In a forward genetic screen aimed at identifying Rabs that regulate phagocytic processing of neuronal mHTT_ex1_ aggregates, we uncovered glial Rab10 as a modifier of prion-like spreading of mHTT_ex1_ from neurons to glia. Interestingly, several Rab proteins have been previously reported to alter secretion or cell-to-cell propagation of tau or α-synuclein aggregates (Rodriguez et al., 2017; Bae et al., 2018; Ugbode et al., 2019; Rodrigues et al., 2022), suggesting that Rab-dependent endomembrane fusion could be exploited by prion-like aggregates. Our data suggest that Rab10 acts downstream of Draper to enable engulfed mHTT_ex1_ aggregates to evade lysosomal degradation, perhaps escaping to the cytoplasm during Rab10-dependent vesicle fusion.

While relatively understudied compared with other Rabs, Rab10 has been reported to localize to and regulate maturing endo/phagosomes and lysosomes in human macrophages and microglia (Cardoso et al., 2010; Lee et al., 2020). Interestingly, Rab10 has already been implicated in multiple neurodegenerative diseases: (1) *RAB10* gene expression is altered in human AD brains (Ridge et al., 2017), (2) rare *RAB10* polymorphisms are linked to protection against AD (Ridge et al., 2017; Tavana et al., 2018), (3) Rab10 depletion is associated with reduced Aý levels (Udayar et al., 2013; Ridge et al., 2017), and (4) Rab10 is a substrate of the PD risk factor gene leucine-rich repeat kinase 2 (LRRK2) (Steger et al., 2016; Seol et al., 2019). Mutant LRRK2-mediated Rab phosphorylation leads to endocytic defects (Liu et al., 2020; Rivero-Ríos et al., 2020; Streubel-Gallasch et al., 2021), and LRRK2-modified Rab10 localizes to and promotes secretion from stressed lysosomes (Eguchi et al., 2018; Kluss et al., 2022). Interestingly, phosphorylated Rab10 (pRab10) levels are elevated in the CNS of AD and PD patients, and pRab10 has been reported to colocalize with pathological tau (Yan et al., 2018; Tezuka et al., 2022) suggesting that post-translational modification of Rab10 modifies the disease state. Whether Rab10’s phosphorylation status affects its ability to regulate prion-like activity of mHTT_ex1_ or other amyloid aggregates warrants further investigation.

In conclusion, our findings demonstrate that axonal mHTT_ex1_ aggregates activate glial phagocytosis but also impair normal glial responses to acute neural injury. Aggregate engulfment-induced defects in endolysosomal processing, such as Rab-mediated vesicle membrane fusion, could facilitate the formation and spread of prion-like aggregate seeds in the brain. These findings point to central roles for phagocytic glia in regulating pathological aggregate burden in HD and highlight the importance of exploring therapeutic interventions that target non-neuronal cells.

## ACKNOWLEDGMENTS

We would like to thank Sabrina Abbruzzese, Ryan Buist, Georgiana Moore, and David Tomlinson for experimental assistance, and all former and current members of the Pearce Lab for their support and many fruitful discussions about this project. This work was supported by funding from University of the Sciences, Rowan University, the W.W. Smith Charitable Trusts, and the National Institutes of Health (awards NS128847 and AG063295).

## TABLE LEGENDS

Table 4. Statistical information for all quantitative results.

## Notes

### Competing Interest Statement

The authors have declared no competing interest.

### Summary of Updates

Figures reorganized and new data added.

